# Hydrogen-bonding changes cause differences in imipenem breakdown activity in OXA-48 variants

**DOI:** 10.64898/2026.01.20.700306

**Authors:** Daojiong Wang, Adrian J. Mulholland, James Spencer, Marc W. van der Kamp

## Abstract

The *β*-lactamase OXA-48 efficiently hydrolyses carbapenem antibiotics, especially imipenem. Carbapenem resistance is a rising clinical concern, and is frequently associated with OXA-48 and its variants. OXA-48 variants carrying different mutations in the *β*_5_-*β*_6_ loop differ in hydrolytic activity towards imipenem. OXA-517 has a higher *K*_M_, but similar *k*_cat_ for imipenem hydrolysis, compared to OXA-48, whereas OXA-163 and -405, which have similar mutations in the *β*_5_-*β*_6_ loop, are less active. Multiscale simulations (using quantum mechanics/molecular mechanics, QM/MM) of deacylation of the respective imipenem acylenzymes show this to be most efficient when the deacylating water (DW) acts as a hydrogen bond (H-bond) donor to imipenem, and the carboxylated Lys73 base is less hydrated. Calculated barriers for deacylation correlate very well with experimental data, but for OXA-163 and -405 only when DW acts as a H-bond acceptor. Dynamics simulations of imipenem acylenzyme complexes show that mutations in the *β*_5_-*β*_6_ loop change the active site H-bond network. In OXA-48, the DW H-bonding pattern linked to high activity is more frequently sampled, and in OXA-517 it is stabilised through H-bonding to Thr213; explaining the higher *k*_cat_ values compared to OXA-163 and -405, where this is not the case. Furthermore, simulations of non-covalent imipenem complexes indicate that increased *K*_M_ for OXA-517 is linked to lower binding affinity, caused by repositioning of bound imipenem. Our work identifies the molecular basis for differences in imipenem hydrolytic activity between OXA-48 variants, offering detailed insights into how active site interactions alter the dynamics and reaction efficiencies related to antibiotic resistance.

## Introduction

Antimicrobial resistance (AMR) has become a serious threat to public health ^1, 2^. The discovery of antibiotics is one of the most important findings in the twentieth century. However, under the pressure to survive, bacteria have evolved or acquired various ways to protect themselves against antimicrobial agents, leading to 1.14 million deaths in 2021 directly attributable to antimicrobial resistance ^3^. One of the major resistance mechanisms in bacteria is the production of *β*-lactamases, enzymes that hydrolyse *β*-lactam antibiotics, the most widely used treatment for bacterial infections worldwide ^4, 5^.

In *β*-lactam antibiotics, the *β*-lactam ring mimics the D-Ala-D-Ala moiety of the natural substrate of penicillin-binding proteins (PBPs), a family of enzymes essential to maintaining bacterial cell wall structure and stability ^6, 7^. Bacteria can produce enzymes called *β*-lactamases (BLs) that hydrolyse the *β*-lactam ring, thus leading to antibiotic inactivation ^8^. BLs are divided into four classes, A, C, D and B, based on the Ambler (sequence-based) classification ^9^ system: Class B BLs are zinc-dependent, whereas classes A, C and D are serine BLs that all have an active site serine residue that performs a nucleophilic attack on the *β*-lactam ring, and so are mechanistically related to the transpeptidase activity of PBPs ^10, 11^. The OXA-48-like family of class D *β*-lactamases is particularly relevant for antibiotic resistance, due to their wide distribution ^12^ and the fact that many members of the family have significant hydrolytic activity against carbapenems, which used to be considered ‘last-resort’ antibiotics ^13^. The OXA-48 parent enzyme has a higher hydrolytic activity towards imipenem than other carbapenems ^14^. The presence of OXA-48-like enzymes, in combination with other resistance mechanisms, can lead to high levels of carbapenem resistance ^13, 15^.

The general mechanism of imipenem hydrolysis by OXA-48-like enzymes consists of three main stages: initial non-covalent binding (Michaelis complex formation), acylation and deacylation (**Figure 1**) ^14^. Both acylation and deacylation involve a general base. Unlike class A BLs, which are expected to utilize Glu166 as the general base in both acylation and deacylation ^16–18^, class D BLs such as the OXA-48-like enzymes employ a carboxylated Lys73 for both ^19, 20^. After the formation of the non-covalent Michaelis complex, acylation occurs, in which the carboxylated Lys73 (KCX) functions as a general base to activate Ser70 ^19^. Ser70 then performs a nucleophilic attack upon the carbonyl carbon of the *β*-lactam ring, opening the ring and forming a covalent bond with the electrophilic carbon. The resulting acylenzyme complex (AC) is deacylated to release the product and let the enzyme turn over: this step was determined to be the rate-limiting step in imipenem hydrolysis by OXA-48 and -163 (through pre-steady state kinetic analysis) ^21^. Deacylation involves an active site water molecule, known as the deacylating water (DW), which is activated by KCX to act as a nucleophile. This nucleophilic attack on the carbonyl carbon leads to a short-lived tetrahedral intermediate (TI), with the energy barrier associated with this step explaining the difference between OXA-48 activity towards imipenem and meropenem ^22^, further suggesting that the TI formation in deacylation is rate-determining. The second step of deacylation involves bond breakage between Ser70 and hydrolysed imipenem, followed by release of the final product. The expectation is that the imipenem pyrroline ring is in its Δ^2^ (enamine) tautomeric form to allow for efficient deacylation, as was confirmed for the Class A carbapenemase KPC-2 ^23^. As well as the Δ^2^ tautomer, both stereoisomers of the alternative Δ^1^ tautomer (*R*-Δ^1^ and *S*-Δ^1^, imine), have been observed in OXA-48 acylenzyme crystal structures ^21, 24–26^. However, the Δ^1^ tautomer may inhibit the enzyme ^27^, consistent with NMR studies that indicate OXA-48 and other carbapenemases form a Δ^2^ or an *R*-Δ^1^ deacylation product (with the latter potentially occurring through rapid non-enzymatic tautomerisation) ^28^.

**Figure 1.**
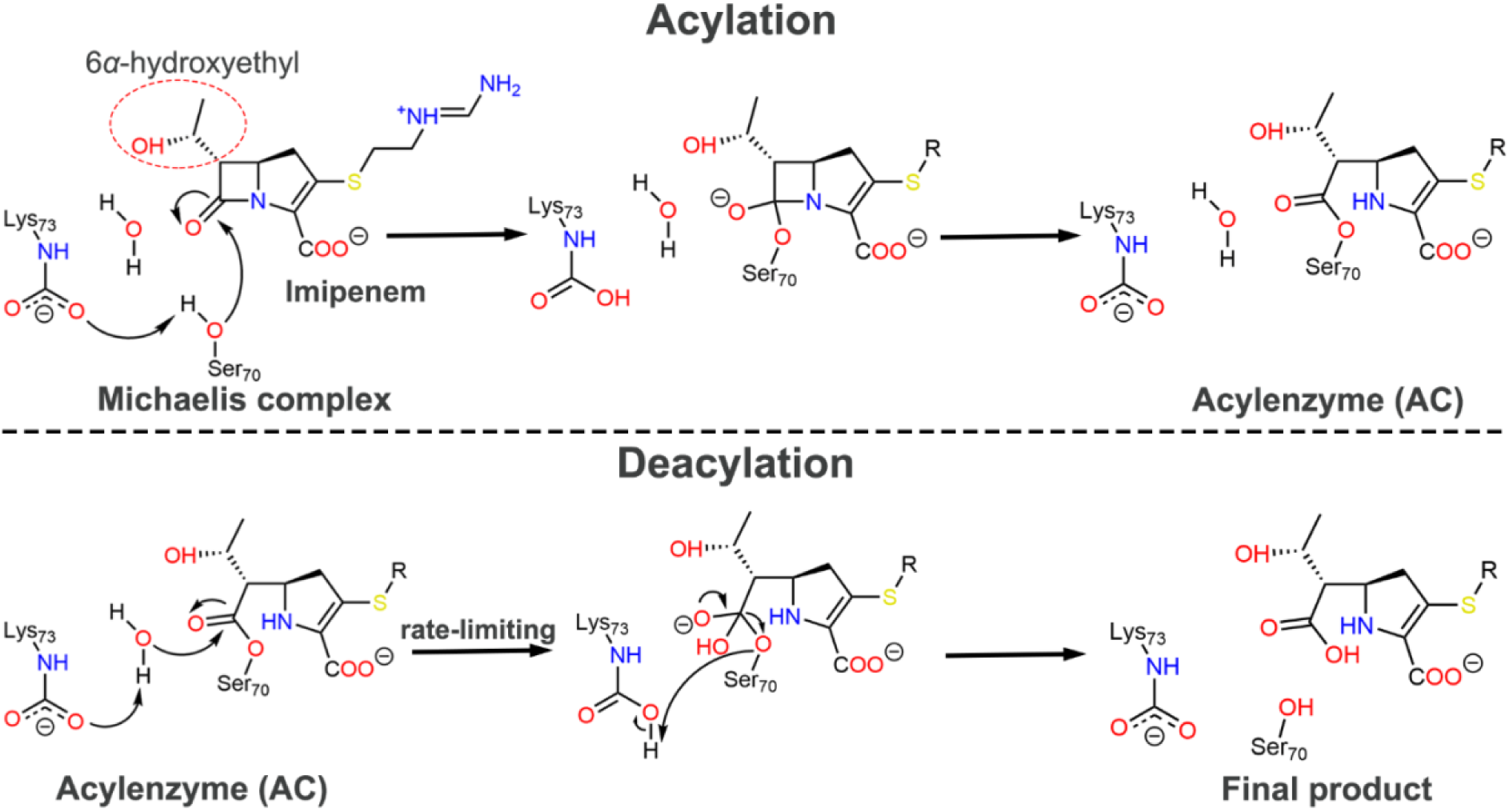
Reaction mechanism of imipenem hydrolysis (via its Δ^2^ tautomer) by OXA-48-like enzymes ^10, 14^. Carboxylation of Lys73 is required for efficient hydrolysis ^19, 20^.

OXA-48-like enzymes (as well as most other class D BLs) have three widely conserved active site motifs ^14, 20^. Motif I (S^70^-X-X-K^73^) contains Ser70 and Lys73, which can be carboxylated, both essential for the reaction (see above). Although motif II (S^118–^V-V) and motif III (K_208–_T-G) are distant from motif I in sequence, they are located close to the key catalytic residues (on a helix and a beta-strand, respectively). Val120 forms a “water channel” with Leu158, allowing a water molecule (DW) to enter the active site and support deacylation ^24, 29^. Val120 also provides a hydrophobic environment that lowers the p*K*_a_ of Lys73, thereby facilitating its carboxylation ^30, 31^. Lys208 from motif III forms a hydrogen bond to the sidechain of Ser118 and has been proposed to be involved in substrate recognition ^31^. Alanine substitution at Ser118 and Lys208 was reported to cause a significant reduction in MICs for carbapenems ^32^. The Ω loop (residues Tyr144 to Arg163 in OXA-48) and *β*_5_-*β*_6_ loop (residues Thr213 to Lys218 in OXA-48) are not directly involved in reaction, but these loops have been indicated to play important roles in enzyme specificity. The enhanced flexibility of the Ω and *β*_5_-*β*_6_ loops in lab-evolved OXA-48 variants (compared to the parent OXA-48) has been suggested to cause increased efficiency towards ceftazidime ^33–35^.

Several natural, clinically relevant, variants of OXA-48 have changes in the *β*_5_-*β*_6_ loop. OXA-163, a commonly encountered OXA-48 variant, contains a four-residue deletion (214-RIEP-217) and single residue substitution (S212D). OXA-163 possesses significant cephalosporin hydrolysis activity ^36^, whereas carbapenem hydrolysis efficiency, particularly for imipenem, is significantly reduced compared to OXA-48 (**Figure 2**) ^21, 37^. It was hypothesised, based on single MD trajectories of substrate complexes, that this reduction is related to multiple conformations or substates being sampled by OXA-163 (compared to OXA-48) ^21^. Another variant, OXA-405, has a similar four-residue deletion (213-TRIE-216), leading to similar effects on hydrolytic activity. The replacement of the *β*_5_-*β*_6_ loop of OXA-48 with that of OXA-18, which can hydrolyse cephalosporins, also results in the ability to hydrolyse cephalosporins ^38^. In both of these variants Arg214 in the *β*_5_-*β*_6_ loop, which can form a salt bridge interaction with Asp159 to help stabilise the Ω loop in OXA-48 resulting in a higher melting point of the protein, is missing ^39^. Mutation studies of Arg214 in OXA-48 and OXA-232 suggest that this residue is crucial for carbapenemase activity by OXA-48-like enzymes ^40^.

**Figure 2.**
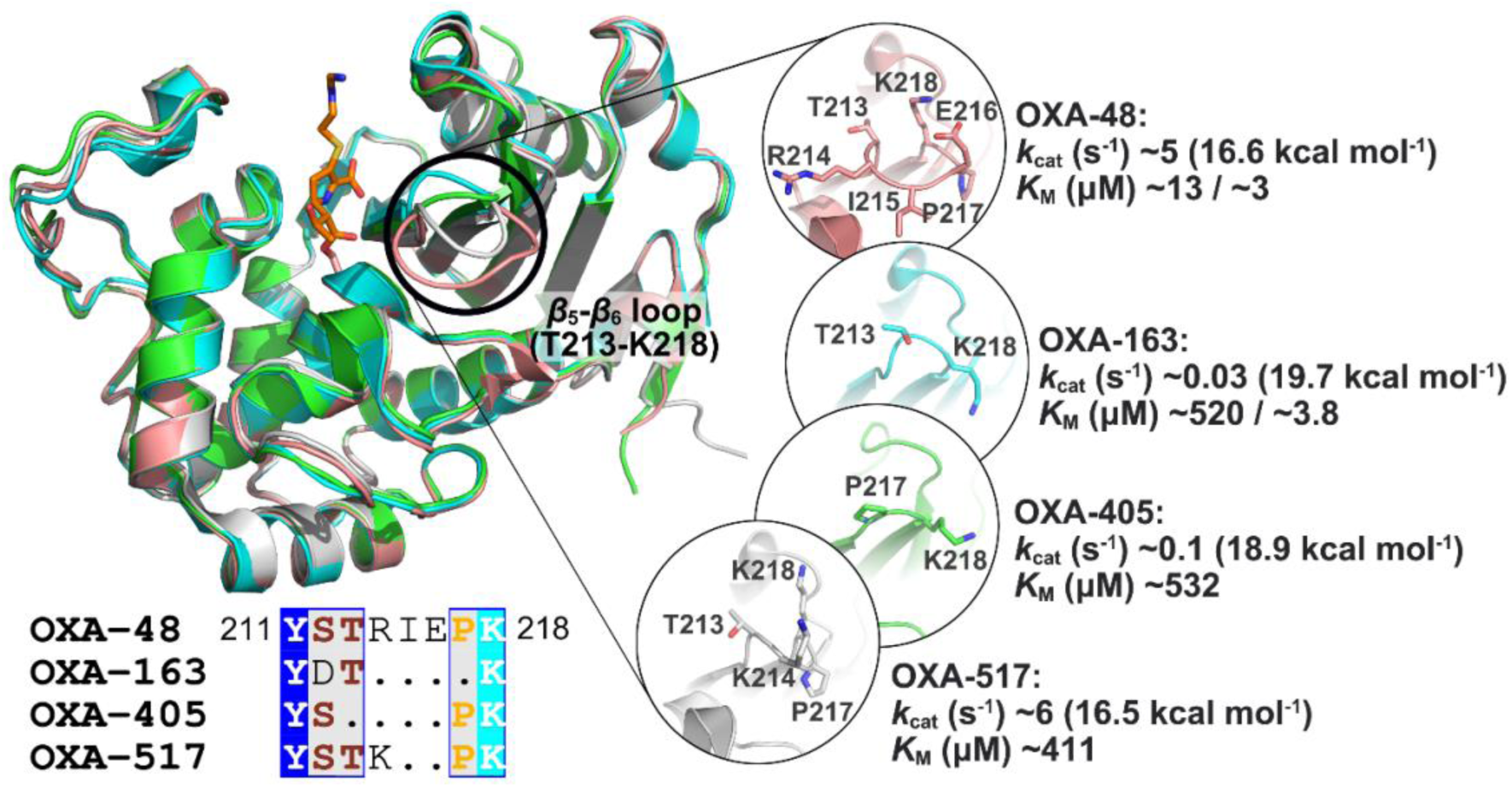
Aligned crystal structures of OXA-48 (pink, PDB ID: 6P97 ^24^, chain A, imipenem acylenzyme complex), OXA-163 (cyan, PDB ID: 7KHZ ^21^, chain A, imipenem acylenzyme complex), OXA-405 (green, PDB ID: 5FDH ^42^, chain A, apo) and OXA-517 (grey, PDB ID: 6HB8 ^41^, chain A, apo) (upper panel). For clarity, the covalently bound imipenem is shown as sticks (orange carbons) only in OXA-48. The position of the imipenem core in OXA-163 is highly similar. Sequences of the *β*_5_-*β*_6_ loop are displayed below (full sequence alignment in **Figure S1**). Structural differences in the *β*_5_-*β*_6_ loops are highlighted in the inserts, together with experimental kinetic data obtained from published studies ^20, 21, 37, 41, 42^. Different experimental *K*_M_ values of OXA-48 and -163 have been reported, as indicated. Experimental *k*_cat_ values were converted into activation free energies (Δ^‡^*G*, in kcal mol^-1^) at 300 K using transition state theory ^43^.

Recently, a new OXA-48 variant, OXA-517, was discovered ^41^. OXA-517 harbours a two-residue deletion (215-IE-217) and one residue substitution (R214K) in the *β*_5_-*β*_6_ loop, compared to OXA-48. Notably, even though OXA-517 has mutations in the *β*_5_-*β*_6_ loop, it shares a similar *k*_cat_ value for imipenem hydrolysis with OXA-48 (whereas *k*_cat_ values are much lower for OXA-163 and -405; **Figure 2, Table S1**) ^20, 37, 41, 42^. However, OXA-517 does have a higher *K*_M_ value than OXA-48, causing a lower hydrolysis efficiency overall. Clearly, the main difference between OXA-48 and the variants mentioned (OXA-163, -405 and -517) is in the *β*_5_-*β*_6_ loop (**Figure 2 and S1**), but how mutations in the *β*_5_-*β*_6_ loop affect carbapenemase activity remains unclear.

Here, we use detailed atomistic simulations to investigate the differences in imipenem hydrolysis observed for OXA-48-like enzymes with changes in their *β*_5_-*β*_6_ loops. Multiscale (QM/MM) reaction simulations were used to investigate the rate-limiting deacylation step. Our results support that for each variant studied, the energetically most favourable situation for deacylation is a less hydrated catalytic base (KCX), with the DW acting as hydrogen bond donor to the imipenem 6*α*-hydroxyethyl group, as previously found for OXA-48 ^22^. Molecular dynamics simulations and QM/MM calculations of the different acylenzyme complexes revealed a significant difference between the variants in the preference of H-bonding patterns formed around the DW. Whereas the differences in the *β*_5_-*β*_6_ loop disfavour the DW acting as hydrogen bond donor in OXA-163 and -405, OXA-517 still allows this interaction, which favours imipenem deacylation. Furthermore, molecular dynamics simulations of the non-covalent imipenem complexes show that in OXA-517, the imipenem position differs from OXA-48, leading to lower binding affinity and thus a change in *K*_M_. Overall, our results explain the changes in hydrolytic activity towards imipenem for key variants of OXA-48 with mutations in the *β*_5_-*β*_6_ loop, providing information on how subtle conformational differences cause changes in enzyme activity, aiding in a deeper understanding of antibiotic resistance conferred by *β*-lactamases.

## Methods

### System setup and preparation

Simulations applied protocols similar to those used in previous studies ^22, 44^. The X-ray crystal structures of imipenem acylenzyme complexes binding with OXA-48 and OXA-163 were obtained from the PDB database (PDB ID: 6P97 ^24^, chain A, and PDB ID: 7KHZ ^21^, chain A, respectively). The initial structures of imipenem acylenzyme complexes of OXA-405 and OXA-517 were generated by modelling imipenem into the apo X-ray structures of OXA-405 (PDB ID: 5FDH ^42^, chain A) and OXA-517 (PDB ID: 6HB8 ^41^, chain A) through alignment with the OXA-163 and OXA-48 acylenzyme complexes, respectively. The missing atoms of the carboxylated Lys73 were added to OXA-48, -405 and -517 acylenzyme complexes, and the deacylating water (DW) was added manually to all. The OXA-163 acylenzyme complex, in which Lys73 is substituted by Ala73 in the crystal structure, was reverted to a carboxylated Lys73. Non-covalent (Michaelis) complexes of imipenem with the enzymes were generated through breaking the bond between Ser70 and imipenem in the previously prepared acylenzyme complexes and reforming the *β*-lactam ring manually in PyMOL. Partial charges for the acylated imipenem (including Ser70) were calculated by using RESP charge calculation based on HF/6-31G(d) of the capped Ser70-imipenem fragment using the R.E.D. server ^45^. For the non-covalently bound imipenem, partial charges were determined using the AM1-BCC model using antechamber ^46^. The GAFF2 force field ^47^ was used to describe the atom types and bonded parameters of both forms of imipenem. All waters in the crystal structures were retained (but ions and buffer molecules deleted) and the protonation states of ionizable residues at pH 7.0 were determined using PropKa3.1 ^48^, with histidine tautomers predicted with the reduce program from AmberTools ^49^. All ionizable residues were predicted to be in their standard protonation states (Asp and Glu deprotonated; Lys and Arg protonated), with His34, His90, His109, His178 and His182 singly protonated on NE2, His38 and His140 singly protonated on ND1. Protein atoms were treated with the Amber ff14SB force field ^50^. All complexes were solvated in a rectangular box of TIP3P water ^51^, with a minimum distance between the protein and the box edge of 10 Å. To neutralise the complexes, one Na⁺ ion was added to OXA-48 and OXA-405 and two Na⁺ ions were added to OXA-163, by replacing random bulk water molecules (no ions were added to OXA-517).

### MM molecular dynamics (MD) simulation and analysis

All systems were first minimized for 2000 cycles (1000 cycles with steepest descent followed by 1000 cycles with conjugate gradient). Then, the temperature was increased from 50 to 300 K over a period of 20 ps. During all equilibration and production simulations, periodic boundary conditions were applied and the SHAKE algorithm was applied to fix all bond lengths involving hydrogen atoms. A time step of 2 fs was used and cutoff radius for non-bonded interactions was set to 8.0 Å.

For each acylenzyme complex, five independent simulations were run. Initially, unrestrained simulations of 120 ns were run in the NPT ensemble, at 300 K (maintained using Langevin dynamics, collision frequency 0.2 ps^−1^) and 1 bar (using Berendsen barostat with isotropic position scaling, pressure relaxation time 1 ps). The carbapenem 6*α*-hydroxyethyl group can adopt three rotameric states or orientations (according to dihedral angle, C7-C6-C16-O7) (**Figure S2**) ^52, 53^, and previous MM MD simulation of the OXA-48 imipenem acylenzyme complex shows that the 6*α*-hydroxyethyl group can adopt all three ^22^. QM/MM simulation of imipenem (and meropenem) hydrolysis by OXA-48 indicates that the different orientations significantly influence the energy barrier, with orientation I (∼50°) the most reactive rotameric state ^22^. Therefore, this orientation was enforced here. First, one frame per replica was selected from the initial restraint-free production simulations, in which the imipenem 6*α*-hydroxyethyl group adopted the rotamer captured in the OXA-48 and OXA-163 imipenem acylenzyme crystal structures. Then, the imipenem 6*α*-hydroxyethyl group was gradually changed to ∼50°, by applying a dihedral angle restraint (force constant of 100 kcal mol^−1^ rad^-2^) and changing it by 5° every 100 ps (27 steps). The final frame from the last step was used to perform a further 120 ns MM MD simulation with a two-sided harmonic dihedral angle restraint (force constant of 100 kcal mol^−1^ rad^−2^) on the 6*α*-hydroxyethyl group, allowing restraint-free sampling from 0° to 100°. The first 20 ns of this simulation was regarded as further (post-restraint) equilibration and was excluded from trajectory analysis. The trajectories were saved every 10 ps, resulting in 10000 frames per replica for analysis; a total of 50000 frames per system.

To investigate differences in relative binding energy, ten independent simulations of Michaelis complexes (imipenem non-covalently bound) were conducted of 10.5 ns each (with the first 0.5 ns simulation treated as equilibration). Because imipenem can (partially) dissociate from enzyme in simulation, two weak one-sided harmonic restraints were applied throughout the simulations for the oxyanion hole hydrogen-bond distances, to capture the acylation-ready state.

The restraints came into force when the distance between the carbonyl oxygen of imipenem and the backbone amide H atom of Ser70/Tyr211 was above 2.0 Å (force constant 10 kcal mol^−1^ Å^−2^). The trajectories were saved every 10 ps, resulting in 1000 frames per replica and a total of 10000 frames per system were used for analysis.

All simulations were conducted using Amber20/AmberTools20 and trajectories were analysed using CPPTRAJ ^49, 54^. Heavy atom RMSD values of the Ω loop were calculated by aligning trajectories on all C*α* atoms, excluding those in the Ω loop. The first frame of each trajectory was used as reference. Differences in C*α* fluctuation (ΔRMSF_C*α*_) were calculated by using RMSF per residue of each variant minus the RMSF value of corresponding residue in OXA-48. The RMSF values for each residue C*α* were determined by first obtaining the average coordinate set of each replica and then aligning the trajectory to this coordinate set prior to RMSF calculation. An independent two-sample, two-tailed t-test was performed to compare the per-residue RMSF values of OXA-48 variants with those of WT OXA-48 (n=5). Clustering analysis was conducted with the k-means algorithm, and the number of clusters was determined based on the highest pseudo-F statistic (pSF) value. For clustering of acylenzyme complexes, the trajectories were first aligned to global average structure (residues Arg214-Pro217 in OXA-48 and Lys214-Pro215 in OXA-517 were excluded to ensure positional correspondence across all enzymes), after which clustering was performed based on RMSD of C*α* atoms. For clustering of Michaelis complexes, the trajectories were first aligned to the C*α* atoms of active site residues that are located within 5 Å of the ligand (residues Ile102-Trp105, Lys116-Val120, Lys208-Ser212, Leu247-Arg250 in OXA-48). Then, clustering was performed based on the RMSD of heavy atoms of the imipenem core (**Figure S3**). Principal component analysis was performed after aligning trajectories to the global average structure. For hydrogen bond analysis, the default criteria in CPPTRAJ were used (a hydrogen donor-acceptor distance less than 3.0 Å, and a donor-hydrogen-acceptor angle between 135° and 180°).

### QM/MM MD reaction simulation and analysis

To investigate the reaction barrier of deacylation, 2D QM/MM umbrella sampling MD was applied to the acylenzyme complexes of imipenem with OXA-48-like proteins. Frames from last 40 ns of MM MD simulation were selected based on a reactive DW position (distance between DW@O and KCX@OQ1 closer than 3.0 Å and distance between DW@O and electrophilic carbon closer than 3.5 Å), together with the desired DW H-bonding pattern and hydration state. The core structure of imipenem, sidechains of Ser70 and carboxylated Lys73, and the deacylating water are included in the QM region (**Figure S3**). The QM region comprises 43 atoms (including 3 link atoms) and carries a total charge of –2 e. The QM region was described by the semi-empirical method DFTB2 (SCC-DFTB) ^55^. DFTB2 was shown to provide reasonable geometries and to reproduce experimental differences in enzyme kinetics for carbapenem deacylation in OXA-48 in our previous work ^22^. Simulation settings for QM/MM were identical to the previous MM simulations, apart from using a 1 fs timestep (and no SHAKE applied on the QM region). Selected frames were used to calculate a free energy surface (FES) using umbrella sampling (US), applying two reaction coordinates. One is proton transfer coordinate between KCX and DW: distance (*d*[KCX@OQ1, DW@H] – *d*[DW@O, DW@H]) which shows the break of O-H bond in DW and formation of KCX@OQ1-DW@H bond. The other is nucleophilic attack involves with the OH^−^ of DW acting nucleophilic attack (*d*[DW@O, IME@C7]) to electrophilic carbon and forming a covalent bond. From acylenzyme complex to tetrahedral intermediate, the proton transfer coordinate value changes from 0.8 Å to –1.0 Å, while for nucleophilic attack, it changes from 3.5 Å to 1.5 Å (both in steps of 0.1 Å).

Umbrella sampling was first performed window-by-window along the approximate minimum free energy path used in previous work ^22^, then the full 2D surface was covered by going outwards from this minimum free energy path. 399 US windows were used in total, using 2 ps MD sampling in each window for each replica. This allows for sufficient sampling to uncover trends while maintaining the specific active site conformation ^22^. To avoid changes in hydrogen bonding of the waters surrounding the carboxylated Lys73 during the reaction simulations, distance and angle restraints were applied (see Supplementary Note S1 for details). Reaction coordinate values were saved every step and analysed using the Weighted Histogram Analysis Method (WHAM) ^56^. The Minimum Energy Path Surface Analysis (MEPSA) program ^57^ was used to plot the resulting FES and the minimum free energy path.

### QM/MM potential energy calculations for H-bond network changes

To investigate whether the wider H-bond network can support the reaction favoured H-bond pattern of the DW (as hydrogen donor), a series of QM/MM minimisation and single point energy calculations was conducted. Three representative frames from the acylenzyme MD simulation of each system where the DW acts as hydrogen acceptor and Thr213 (if present) is involved in the H-bond network were selected. The core structure of imipenem, sidechain of Ser70, part of Tyr211 backbone (cut at C*α*-C bond), whole Ser/Asp212 backbone and part of Thr213/Pro217 backbone (cut at C*α*-C bond) with the entire sidechain, DW and waters which involved in the H-bond network were included in the QM region (described by DFTB2). For OXA-48, -163 and -405, the QM regions contain 60 atoms (including 5 link atoms), whereas for OXA-517 it contains 63 atoms (including 5 link atoms), due to an additional water molecule. The total charge of each QM region is –1 e. The whole complex was first minimised using the LBFGS ^58^ optimizer, with only atoms within 20 Å of the C*α* atom of Ser/Asp212 allowed to move. The convergence criterion for the energy gradient was set to 0.02 kcal mol^−1^ Å^−1^ and no nonbonded cutoff was applied. The dihedral angle (C6-C16-O7-H8) of imipenem 6*α*-hydroxyethyl hydroxyl group was restrained to –80° (**Figure S2**), using a two-sided harmonic dihedral restraint with a force constant of 3000 kcal mol⁻¹ rad⁻². Then, further minimisations were conducted in a stepwise manner, allowing only atoms within 15 Å of the C*α* atom of Ser/Asp212 to move (to avoid potential energy changes not related to the changing H-bonding pattern). In each step, the dihedral angle was increased by 10° until a local minimum was reached, at which point the deacylating water acted as hydrogen donor and the H-bond network had adjusted correspondingly. The QM/MM potential energy difference was computed between the two local minima, one corresponding to the deacylating water acting as hydrogen acceptor, and the other as hydrogen donor. For accuracy, DFTB2/ff14SB energies were corrected by replacing DFTB2 energies of QM region only with the corresponding M06-2X/def2-TZVP energies, resulting in M06-2X/def2-TZVP//ff14SB energies (with QM-MM interaction terms still at the DFTB2 level). M06-2X/def2-TZVP calculations were performed in ORCA 6.0 ^59^, with the RIJCOSX approximation and the def2/J auxiliary basis set.

### Binding energy calculation (MM/GBSA)

MM/GBSA (with energy decomposition per residue) was performed to calculate the binding energy of imipenem to the different enzyme variants using MMPBSA.py ^60^ from AmberTools20. All trajectory snapshots from the Michaelis complex simulations were used. Topology files for MM/GBSA calculations were prepared by using the mbondi2 radii, as recommended with the Onufriev, Bashford, Case (OBC) generalized Born model used here ^61^. Residues with significant contributions to the binding energy (absolute value of contribution >0.5 kcal mol^−1^) were selected, and a comparison between OXA-48 variants and OXA-48 was constructed.

## Results and discussion

### Difference in imipenem deacylation rates in OXA-48 variants explained by different orientation of the deacylating water

Our previous work on carbapenem deacylation by WT OXA-48 showed that the rotamer of the carbapenem 6*α*-hydroxyethyl group and the hydration state around the carboxylated Lys73 (KCX) are both important factors influencing the free energy barrier ^22^. When the 6*α*-hydroxyethyl group is in the most reactive conformation (rotamer orientation I, dihedral ∼50°), the DW can form two different H-bonding patterns with the carbapenem 6*α*-hydroxyethyl hydroxyl group (**Figure 3, Figure S2**). Here, we perform QM/MM (DFTB2/ff14SB) umbrella sampling reaction simulations with imipenem in this most reactive orientation, to investigate the differences in *k_cat_* between OXA-48 and its *β*_5_-*β*_6_ loop variants OXA-163, OXA-405 and OXA-517. Different H-bonding patterns and hydration states are considered separately (**Figure 3**).

**Figure 3.**
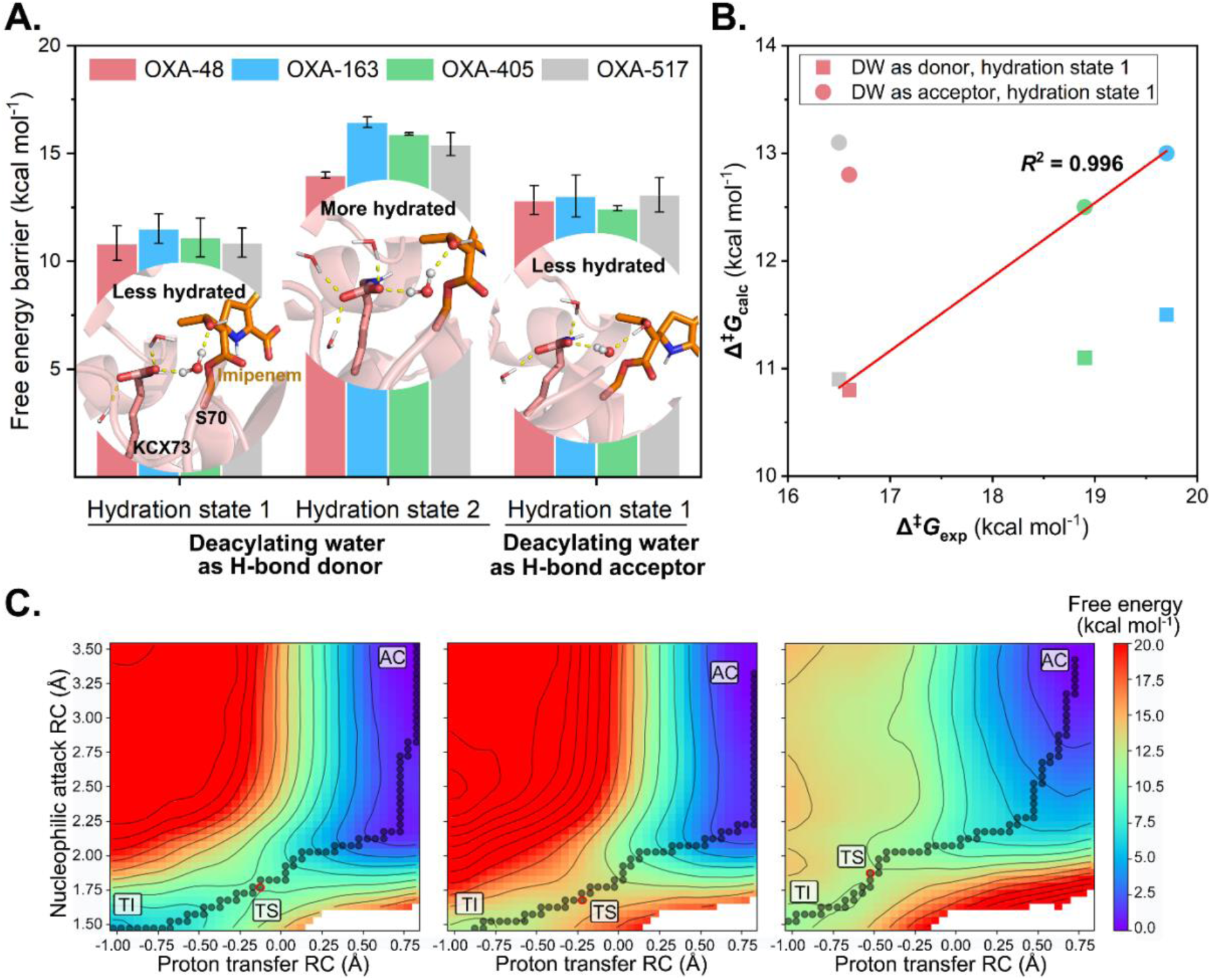
Predicted free energy barriers for deacylation of imipenem acylenzyme complexes of OXA-48 variants in different H-bonding patterns and hydration states. (A) Calculated free energy barriers at the DFTB2/ff14SB level for each state, with error bars obtained from the standard deviation between results of three independent umbrella sampling runs. In the snapshots, the acylenzyme state is shown, with the DW in ball and stick (imipenem with orange carbons). Left: Hydration state 1 (less hydrated, only one water molecules bonded to KCX@OQ2) and DW acts as H-bond donor to the 6*α*-hydroxyethyl group. Middle: Hydration state 2 (more hydrated, two water molecules bonded to KCX@OQ2) and DW acts as hydrogen donor to the 6*α*-hydroxyethyl group. Right: Hydration state 1 and DW acts as hydrogen acceptor to hydroxyl group. (B) Correlation between experimental free energy barriers (values taken from Figure 2) and calculated free energy barriers obtained from umbrella sampling. The same colour scheme as in (A) is used. Correlation is indicated for OXA-48 and -517 with DW acting as H-bond donor, and OXA-163 and -405 with DW as H-bond acceptor. (C) Free energy surfaces for imipenem deacylation by OXA-48, under different hydration states and H-bonding patterns. The order is the same as in panel A. Minimum free energy paths (MFEPs) indicated by graph spheres, with AC = acylenzyme, TS = transition state, TI = tetrahedral intermediate. Similar MFEPs are obtained for the other variants (see **Figure S4**).

For each variant, the highest free energy barrier for deacylation is obtained when KCX is fully hydrated, with both carboxylate oxygens accepting hydrogen bonds from two water molecules. This hydration state has an even larger impact on the barrier for the *β*_5_-*β*_6_ loop variants than for WT OXA-48 (**Figure 3A and Table S2**). When only a single water molecule interacts with the carboxylate oxygen that is not involved in proton transfer (KCX@OQ2), the barriers are lower. When the DW *accepts* an H-bond from the imipenem 6*α*-hydroxyethyl group, the barriers are similar for each variant, and always higher than those obtained with DW *donating* an H-bond to the 6*α*-hydroxyethyl group. In other words, for each variant, the lowest free energy barrier for deacylation is obtained in this situation (DW donating a H-bond to the 6*α*-hydroxyethyl group, KCX@OQ2 accepting a H-bond from a single water molecule). The different H-bonding patterns and hydration states also change the minimum free energy pathway (MFEP), causing a shift of transition state location (**Figure 3C**). In particular, when DW accepts an H-bond from the 6*α*-hydroxyethyl group, the transition state occurs further along the reaction (closer to the tetrahedral intermediate), especially for the proton transfer reaction coordinate. When DW donates an H-bond but KCX is more hydrated, the overall reaction path is very similar, but the TS still occurs slightly later (which is expected, due to a reduction in the KCX proton affinity). These changes in the MFEP are consistent for all four variants studied (**Figures S4** and **S5**), with representative structures of the minima and transition states shown in **Figures S6** and **S7.** Although the order of the predicted energy barriers in the most reactive active site conformation for the four OXA-48-like proteins is consistent with experimental data (10.8 ± 0.8, 11.5 ± 0.7, 11.1 ± 0.9, 10.9 ± 0.7 kcal mol^−1^ for OXA-48, -163, -405, -517 respectively), the differences are too small to explain the experimental *k*_cat_ values ^20, 37, 41, 42^. This suggests that the variation in *k*_cat_ values among these variants may be attributed to differences in attaining or maintaining this most reactive conformation (where a single water molecule interacts with KCX@OQ2 and DW donates an H-bond to the 6*α*-hydroxyethyl group).

For carbapenem hydrolysis by OXA-48 and -163, it has been shown that *k*_cat_ is very close to the deacylation rate: i.e. deacylation is the rate-limiting step ^21^. Owing to the QM method (DFTB2) we used here, our calculated energy barriers for deacylation are consistently underestimated compared to experimental data (**Figure 2**), as found previously for this method for similar reactions ^62, 63^. Our previous work on carbapenem deacylation by OXA-48 demonstrated that DFTB2 underestimates barriers by ∼6.3 kcal mol^−1^ compared to more accurate DFT calculations (M06-2X/def2-TZVP) ^22^. This work also established that the difference in hydrolysis efficiency of imipenem and meropenem by OXA-48 can be attributed to a subtle change in the hydrogen bonding between DW and the 6*α*-hydroxyethyl group, with DW acting as donor with imipenem, but as acceptor with meropenem. Here, if we assume that for both OXA-48 and -517, the DW acts as hydrogen donor with hydration state 1 during the reaction, the barriers, corrected to DFT level (+6.3 kcal mol^−1^), would be 17.1 and 17.2 kcal mol^−1^, respectively. Further, if we assume that for OXA-163 and -405, DW accepts a hydrogen bond from the 6-hydroxyethyl moiety instead, the corrected barriers would be 19.3 and 18.8 kcal mol^−1^. For all four enzymes, these values are close to those inferred from *k*_cat_ values (**Figure 2**), and there is an excellent correlation between the calculated and experimental free energy barriers of the four variants (**Figure 3B**, *R*^2^ > 0.99). These results further support that TI formation in deacylation is (primarily) rate-determining for carbapenem hydrolysis by OXA-48-like enzymes. This conclusion is consistent with previous findings for both ceftazidime and carbapenem hydrolysis by these enzymes ^22, 28^, as well as carbapenem hydrolysis by class A BLs ^62, 63^. In addition, these results indicate that the difference in catalytic rate between OXA-48 and -517, on the one hand, and OXA-163 and -405, on the other, lies in the orientation (and thus hydrogen bond pattern) of the DW. To understand the origin of this subtle difference, we analyse the conformational dynamics of the respective complexes in the following sections.

### β_5_-β_6_ loop mutations cause differences in acylenzyme dynamics around the active site

Molecular dynamics simulations of the acylenzyme states were performed for each OXA-48 variant studied (5 independent simulations of 120 ns for each), to investigate if the variants have differences in conformational preferences and flexibility (that may be correlated to H-bonding patterns causing differences in deacylation). In the simulations, the ‘reactive’ orientation of the 6*α*-hydroxyethyl group was maintained. Combined clustering based on the backbone RMSD (C*α* atoms) shows a clear separation in conformational preference among the variants (**Table S3**). Two distinct clusters are identified, with the following main differences: 1) the backbone distance between Val120 and Leu158, and 2) the conformation of the *β*_5_-*β*_6_ loop (**Figure 4A**). Cluster 1 is dominant in OXA-48 and -517, and has a larger average distance between the C*α* atoms of Val120 and Leu158 (11.1 Å) compared to that observed in cluster 2 (10.4 Å), dominant in OXA-163 and -405 (**Figure S8A**). This points to a difference in the width of the water channel formed by the Val120 and Leu158 sidechains. When measuring the width of the water channel directly (**Figure 4B**), it is clear that OXA-48 always samples a wider water channel, whereas a more closed water channel dominates in OXA-405. If and how this change in water channel width effects deacylation efficiency is not immediately clear. On the one hand, a wider channel may lead to a somewhat higher likelihood of the DW being in a catalytically relevant position in OXA-48 and -517, as indicated by the radial distribution function (RDF) of water molecules around the electrophilic carbon of imipenem and the oxygen atom of KCX that receives the proton (**Figure S8B**, first peaks). However, a reactive DW (based on an ‘ideal’ DW position for deacylation of ≤ 3.0 Å between DW@O and KCX@OQ1 *and* ≤ 3.5 Å between DW@O and the electrophilic carbon of imipenem), was observed only slightly more frequently in OXA-48 (**Table S4**). On the other hand, a wider water channel may lead to a somewhat higher hydration around KCX@OQ2, although the catalytically favourable hydration state 1 is still dominant among all four imipenem acylenzyme complexes (average number of solvent H-bond formed with KCX@OQ2 is 1.45, 1.27, 1.06 and 1.45 for OXA-48, -163, -405 and -517 respectively). Overall, there are some changes in water access to KCX and the electrophilic imipenem between the OXA-48 variants (which was previously suggested based on the more open active site in OXA-163 ^21^), but they are quite subtle and unlikely to lead directly to significant differences in deacylation efficiency.

**Figure 4.**
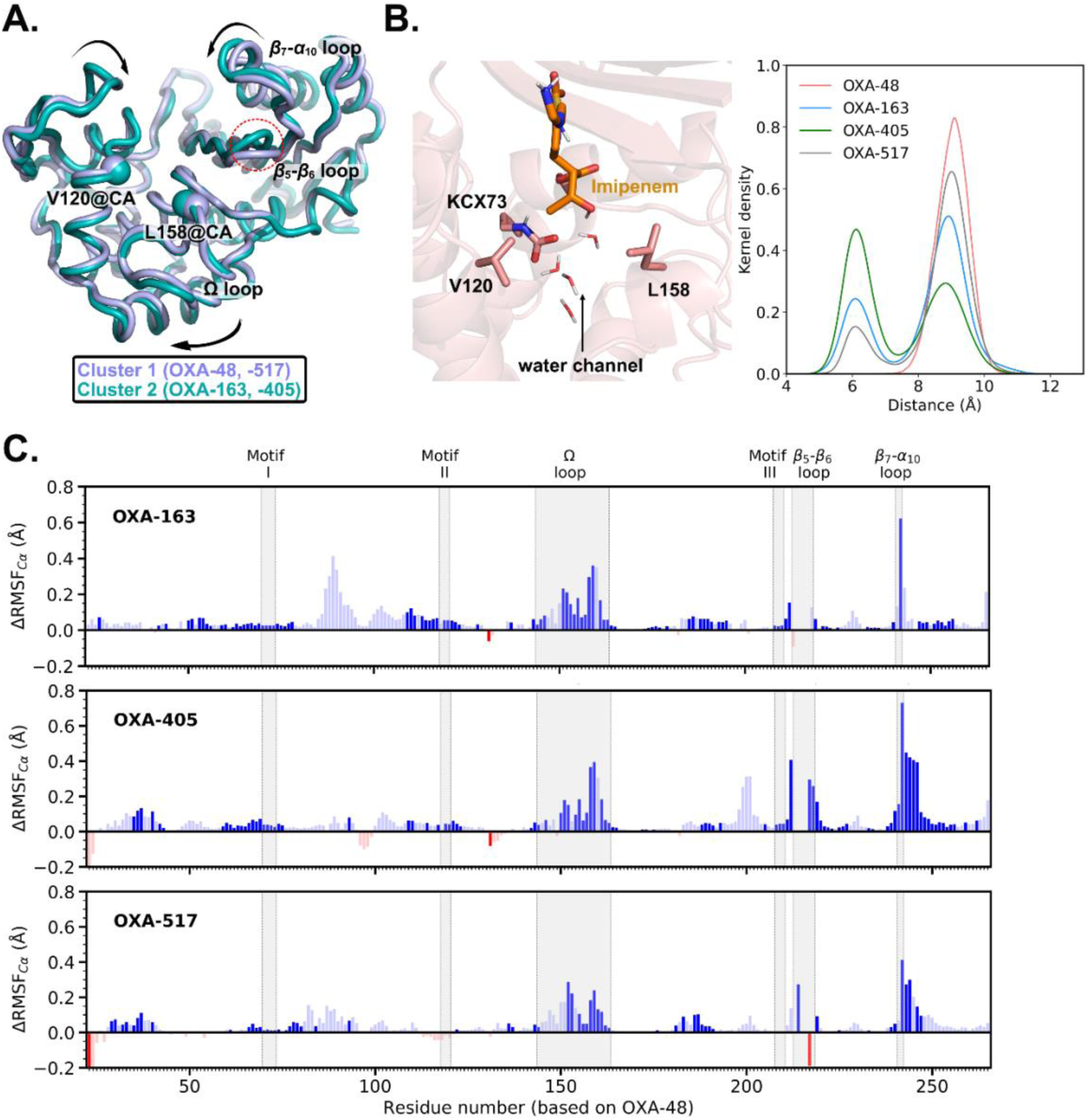
Analysis of imipenem acylenzyme complex dynamics in OXA-48 and the OXA-48 variants. (A) Representative structures of clusters of OXA-48-like proteins based on C*α* atoms, with cluster 1 dominant in OXA-48 and -517, cluster 2 in OXA-163 and -405 (see **Table S3**). (B) Water channel between Val120 and Leu158 in OXA-48 proteins (left) and distributions of the closest distance between Val120 and Leu158 sidechains sampled (right). (C) Change in flexibility (ΔRMSF_C*α*_) of OXA-163, -405 and -517 compared to OXA-48. Increase of RMSF_C*α*_ is coloured in blue, decrease in red. ΔRMSF_C*α*_ values shown in unfaded colours are statistically significant (p <0.05).

The second main difference in the two clusters identified is in the *β*_5_-*β*_6_ loop conformation (as indicated based only on *β*_5_-*β*_6_ loop residues common to all 4 variants). This is not surprising, given the major changes in this loop. The clear distinction in the *β*_5_-*β*_6_ loop backbone conformation between OXA-163 and -405, compared to OXA-48 and -517, is further confirmed by principal component analysis of the common backbone atoms (**Figure S9**). The four-residue deletions in OXA-163 and -405 change the preferred *β*_5_-*β*_6_ loop conformation (cluster 1) such that it is positioned further away from the Ω loop (**Figure 4A**; **Figure S9C)**. Although OXA-517 also harbours a (two-residue) deletion in the *β*_5_-*β*_6_ loop, it still has a similar *β*_5_-*β*_6_ loop conformation as OXA-48. This difference in positioning is related to the presence or absence of a salt-bridge interaction between the *β*_5_-*β*_6_ and Ω loops: Arg214 in OXA-48 forms a stable salt-bridge with Asp159 in the Ω loop (**Figure S10**), with Lys214 in OXA-517 occasionally forming the equivalent interaction, whereas no salt-bridge is detected for OXA-163 and -405. In turn, this salt-bridge affects the flexibility of the Ω loop. All three OXA-48 variants show increased flexibility of the backbone (see ΔRMSF_C*α*_ analysis in **Figures 4C and S11**), with the main increase observed in the Ω loop, and OXA-517 having a slightly more stable Ω loop than OXA-163 and -405. Another difference in flexibility is observed around the *β*_7_-*α*_10_ loop (where there is also a slight difference in conformation between the identified clusters). The exact reason for this change is not clear, but it may be related to weak interactions with the tail group of imipenem (which is highly flexible in all simulations). Overall, the greater flexibility of the active site loops in OXA-163 and -405 observed here may increase the accessibility of the active site, aiding hydrolysis of bulkier substrates such as ceftazidime, as suggested previously ^33^. (Notably, in a structure of the ceftazidime acylenzyme in OXA-48 P68A, there is a lack of interpretable electron density for the Ω loop ^34^).

### Active site H-bond networks in OXA-48 variants differ due to β_5_-β_6_ loop changes

As well as the difference in backbone conformation of the *β*_5_-*β*_6_ loop mentioned above, the variants studied here show significant differences in the sidechains and their interactions in this loop. In particular, Thr213 is involved in hydrogen bonding networks around the acylated imipenem (**Figure 5A**). This *β*_5_-*β*_6_ loop residue is directly adjacent to the mutation/deletion sites in OXA-163 and -517, and is deleted in OXA-405. Three Thr213 rotamers exist (defined by the *χ*_1_ dihedral angle, N-C*α*-C*β*-OG1): gauche-plus (*g*+) (+60°), trans (*t*) (180°) and gauche-minus (*g*−) (−60° or 300°). In our MD simulations, *g*+ is dominant in all three OXA-48-like enzymes possessing Thr213 (OXA-48, -163 and -517, **Figure S12, Figure 5B**). Even though the preferred rotamer is the same in the variants, the interactions with Thr213 can lead to differences in the active site H-bond network. In the OXA-48 imipenem acylenzyme complex, Thr213 predominantly accepts an H-bond from a nearby water (switching to H-bonding with the Ser212 carbonyl when the rare *g*− rotamer is sampled) (**Figure 5A**, left). Due to the two residue deletions, Thr213 in OXA-517 prefers donating a hydrogen bond to a nearby water molecule, and does not form an alternative H-bond with Ser212 (**Figure 5A**, right). For OXA-163, the situation is different: in the preferred *g*+ rotamer, Thr213 predominantly donates an H-bond to the backbone carbonyl of residue 212 (**Figure 5A**, middle), which is the most commonly found conformation here (**Figure 5B**).

**Figure 5.**
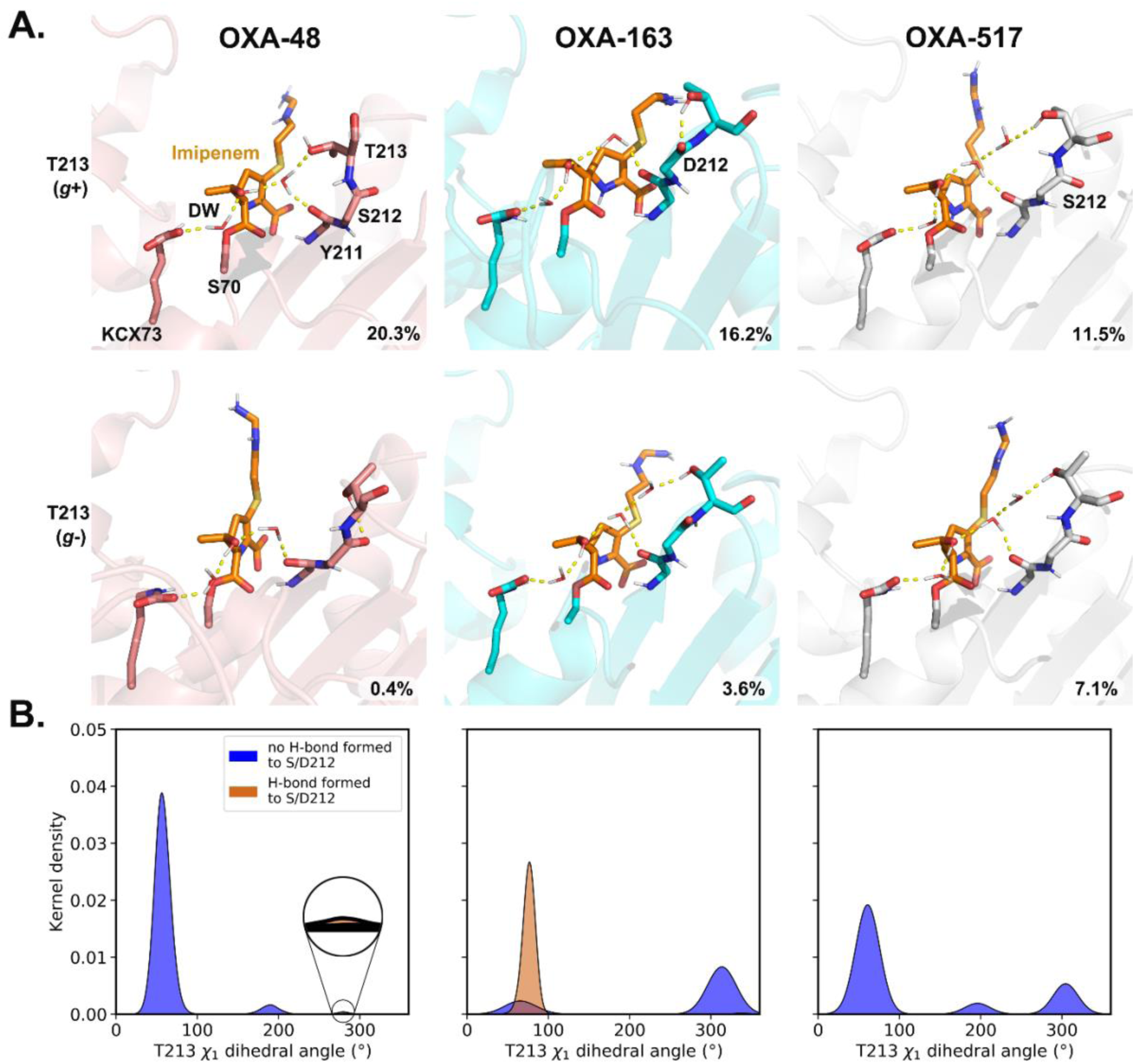
Different H-bond networks among acylenzyme complexes. (A) Representative snapshots of different Thr213 sidechains from simulations. Only polar hydrogens are shown. Key residues (pink) and imipenem (orange) are shown as sticks. H-bonds are shown as yellow dash lines. The fraction of the H-bond network is shown in the bottom right corner; these are relatively low due to the strict criteria (donor-acceptor distance less than 3.0 Å, and a donor-hydrogen-acceptor angle between +/–135° and 180°). (B) H-bond analysis of Thr213 and Ser/Asp212 and analysis of Thr213 sidechain dihedral angle show different H-bonding patterns among acylenzyme complexes (all frames are used here).

In OXA-48, Thr213 is part of a stable H-bond network, where both the sidechain hydroxyl of Thr213 and the backbone carbonyl of Tyr211 act as hydrogen acceptor to a water molecule. This is consistent with what has been observed crystallographically (OXA-48 K73A imipenem acylenzyme complex, PDB ID: 7KH9, chain B) ^21^. This water further accepts an H-bond from the imipenem 6*α*-hydroxyethyl hydroxyl group, leading to the DW donating (rather than accepting) an H-bond to this imipenem hydroxyl (**Figure 5A**, left; **Figure S13**). Because in OXA-163, Thr213 forms an H-bond with Asp212 instead (consistent with PDB ID: 7KHZ, OXA-163 K73A imipenem acylenzyme complex) ^21^, the equivalent water molecule donates H-bonds to both the 6*α*-hydroxyethyl hydroxyl group and the Tyr211 backbone. This then leads to DW acting as hydrogen bond acceptor (**Figure 5A**, middle; **Figure S13**). For OXA-517, both the Thr213 *g*+ and *g*– rotamers donate an H-bond to a nearby water molecule, which in turn interacts with a second water that donates H-bonds to both the Tyr211 backbone carbonyl and the imipenem 6*α*-hydroxyethyl hydroxyl group (**Figure 5A**, right; **Figure S13**). The latter then also leads to DW accepting (rather than donating) an H-bond to the imipenem hydroxyl group. Due to the Thr213 deletion in OXA-405, and thus the lack of a hydrogen bond donor in this position, a similar H-bond network to OXA-163 is typically observed: 45.2% of the frames have a water molecule bridging the 6*α*-hydroxyethyl hydroxyl and the Tyr211 carbonyl, such that the water donates a hydrogen bond to both. (The conformation of the OXA-405 *β*_5_-*β*_6_ loop backbone is very different from the other three OXA-48 variants, see **Figure S9**.) The Ser212-Pro217 peptide bond of OXA-405 is in the *trans* conformation (**Figure S14**), whereas the equivalent E216/K214-Pro217 peptide bond in OXA-48 and -517 adopts the more common *cis* isomer ^64^.)

When only the frames of the acylenzyme MD simulation trajectories are considered that have DW in a ‘reactive’ position (3.0 Å between DW@O and KCX@OQ1 to facilitate proton transfer, and 3.5 Å between DW@O and imipenem carbon to facilitate nucleophilic attack), a significant difference in DW H-bonding preference was found (**Figure 6A**). Specifically, in OXA-48 the DW acts preferentially as a hydrogen donor to the imipenem hydroxyl group, but this is not the case for OXA-163, -405 and -517. As demonstrated above, the DW acting as hydrogen bond donor to the imipenem hydroxyl group favours deacylation. Due to the residue deletion in the *β*_5_-*β*_6_ loop, OXA-517 requires one additional water molecule to bridge the H-bonding interaction between the imipenem hydroxyl group and Thr213 (**Figure 5A**), leading to a preference for the DW acting as acceptor in the acylenzyme MM MD simulations. To better assess the relative energetic difference between the conformations with DW acting either as donor or acceptor to the imipenem hydroxyl group in the OXA-48 variants, we compared QM/MM potential energies of the relevant energy minima, where the hydrogen bond network is treated QM (**Figure 6B** and **Figure S15**). For both OXA-48 and OXA-517, the energy difference is similar and small (2.6 ± 0.6 and 2.2 ± 0.4 kcal mol^−1^, respectively; **Table S5**), indicating that both hydrogen bonding patterns are easily accessible. In contrast, the energy difference is much larger for OXA-163 and -405 (5.3 ± 0.8 and 7.5 ± 1.0 kcal mol^−1^, respectively; **Table S5**), indicating that adopting a reactive conformation with DW as H-bond donor is associated with a significant energy penalty. Overall, these results suggest that in OXA-48 and -517, the water-mediated H-bond network involving Thr213 favours the formation of the DW H-bonding pattern that leads to efficient imipenem hydrolysis.

**Figure 6.**
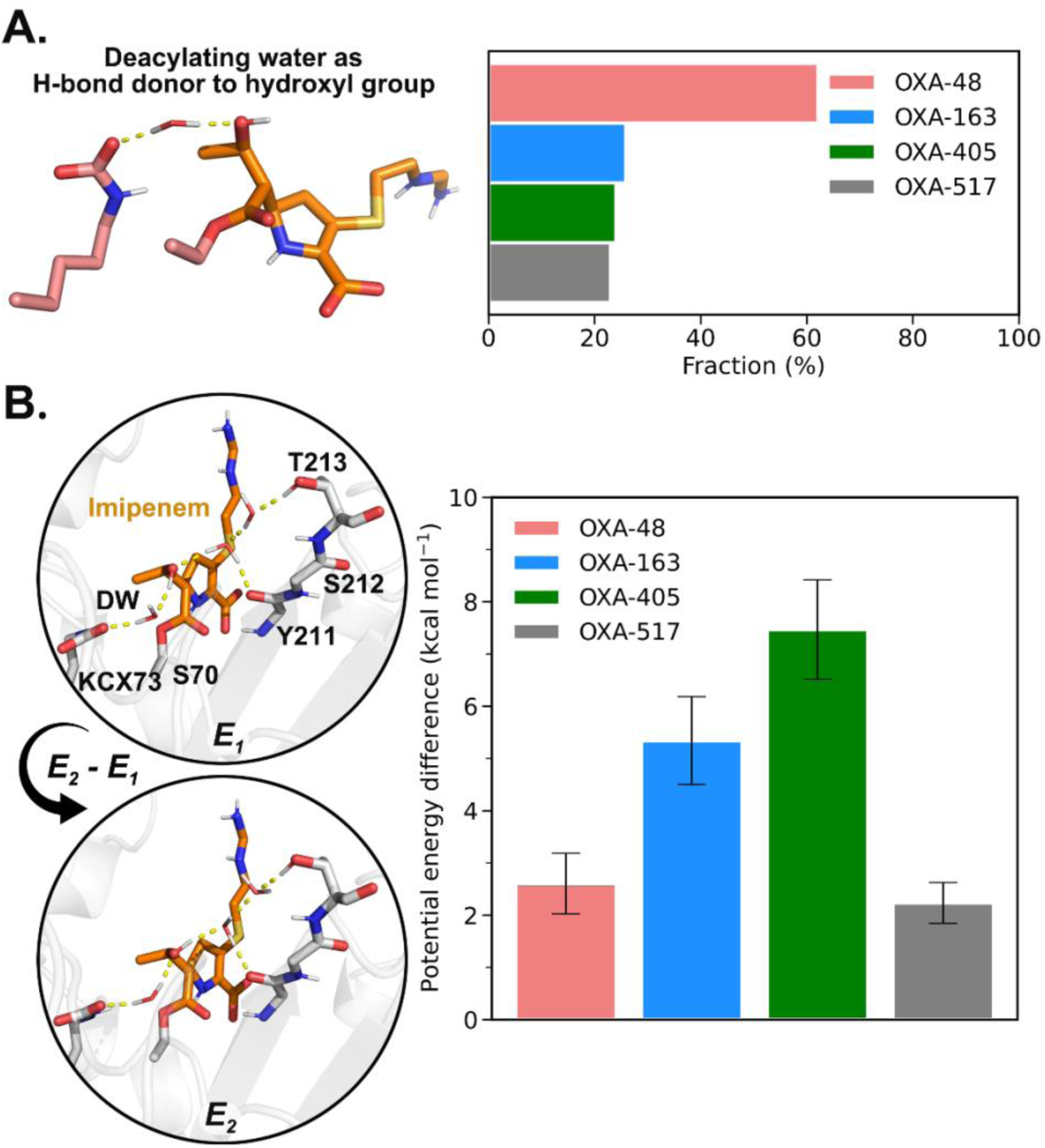
The deacylating water H-bonding pattern that favours imipenem deacylation is energetically preferred in OXA-48 and -517. (A) Analysis of H-bonding patterns between DW and imipenem hydroxyl group of frames with DW in the active site ready for reaction (<=3.0 Å between DW@O and KCX@OQ1 and 3.5 Å between DW@O and the electrophilic carbon). (B) Representative structures at local minima corresponding to different deacylating water H-bonding patterns in OXA-517 (left), used for computing potential energy differences (see **Figure S15** for other variants). Only polar hydrogens are shown. Key residues (grey) and imipenem (orange) are shown as sticks and H-bonds are shown as yellow dash lines. The potential energy differences (*E_2_ – E_1_*) for OXA-48 and its variants (right), calculated at the M06-2X/def2-TZVP//ff14SB level.

### Different imipenem binding pose leads to lower affinity in OXA-517

Although the *k*_cat_ values from experiment indicate similar deacylation rates for OXA-48 and OXA-517, the overall imipenem deacylation efficiency is expected to be significantly lower in OXA-517, due to a >30-fold difference in *K*_M_ (see **Figure 2**). To investigate whether non-covalent binding of imipenem contributes to this difference, we conducted MM MD simulations of Michaelis complexes of imipenem with the OXA-48-like proteins. Clustering of all trajectories based on the carbapenem core of imipenem after alignment on the active site (see Methods) shows a subtle, but significant difference in binding pose between OXA-517 and the other variants (**Figure S16**): imipenem is shifted towards the *β*_5_-*β*_6_ loop.

To estimate if this altered position affects the relative binding affinity, we conducted MM/GBSA analysis on all trajectories. Although this method is approximate, it should be well-suited here for comparison of relative binding energies: the ligand is the same throughout, and the enzyme variants are closely related (i.e. any systematic errors should cancel out). The MM/GBSA calculations indicate that imipenem has a significantly lower binding affinity to OXA-517 than to OXA-48 (p <0.001), whereas the predicted binding energies of imipenem to OXA-48, -163 and -405 are similar (**Table 1**). Decomposing the MM/GBSA binding affinity by residue indicates that the shift in position of bound imipenem in OXA-517 weakens the interactions with Ile102/Trp105/Ser118/Thr209 (compared to OXA-48), resulting in the lower estimated binding affinity (**Figure S17**). Due to the differences in the *β*_5_-*β*_6_ loop (see above), all three variants further show significantly different contributions of *β*_5_-*β*_6_ loop residues to the binding energy, but less favourable contributions from Thr213 (and Pro217) compared to OXA-48 appear to be compensated by preceding residues (Tyr211, and for OXA-163 and -405 also Thr209, Lys208). Our analysis indicates no significant change in overall imipenem binding energy between OXA-48 and OXA-163 (p = 0.27), consistent with no significant difference in *K*_M_ from a direct comparison of kinetics between these variants ^21^. Others also reported *K*_M_ values for OXA-163 and OXA-405 that indicate no significant difference between them ^37, 42^, consistent with the similar binding energies estimated here (p = 0.70). The difference in *K*_M_ values for OXA-163 between refs. ^21^ and ^37, 42^ may thus originate from different experimental protocols. Notably, for these enzymes, where the deacylation rate is particularly slow, the *K*_M_ may also be influenced by accumulation of the acylenzyme (potentially due to tautomerisation of the pyrroline ring to the Δ1 tautomer, that is less reactive ^23^).

**Table 1.**
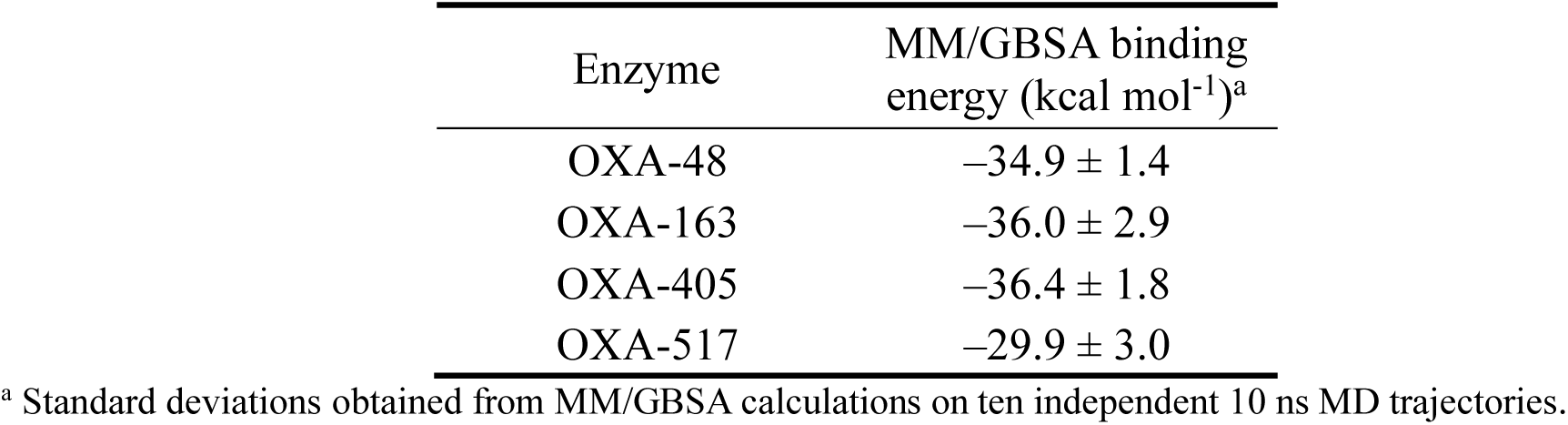
Binding energy of imipenem Michaelis complexes.

| Enzyme | MM/GBSA binding energy (kcal mol<sup>-1</sup>)<sup>a</sup> |
| --- | --- |
| OXA-48 | -34.9 ± 1.4 |
| OXA-163 | -36.0 ± 2.9 |
| OXA-405 | -36.4 ± 1.8 |
| OXA-517 | -29.9 ± 3.0 |

<sup>a</sup> Standard deviations obtained from MM/GBSA calculations on ten independent 10 ns MD trajectories.

## Conclusions

The inactivation of antibiotics is a major contributor to antimicrobial resistance, with the hydrolysis of *β*-lactam antibiotics by *β*-lactamases a primary mechanism ^4, 5, 65^. Understanding the molecular basis of *β*-lactam resistance mediated by *β*-lactamases should help in anticipating and combating antibiotic resistance. Although the enhanced cephalosporin activity by the OXA-163 and -405 variants (compared to OXA-48) may be ascribed to the increased active site volume and flexibility, allowing larger substrates such as ceftazidime to bind and react efficiently, it has remained unclear why these variants also show a significant reduction in carbapenem hydrolysis. For OXA-163 vs. OXA-48, pre-steady-state kinetic studies have shown that this reduction is entirely due to a slower deacylation reaction in OXA-163, but the (atomistic) reasons for this reduction were yet unknown. By conducting detailed simulations of the rate-limiting deacylation reaction, as well as of the acylenzyme complexes, our work here provides such explanations, based on comparison of OXA-48 with three OXA-48 variants with changes in the *β*_5_-*β*_6_ loop: OXA-163, -405 and -517. First, we show that the hydration around the catalytic carboxylated Lys73 (KCX) and the DW H-bonding pattern significantly influences the energy barrier in all variants, as was previously found for OXA-48 ^22^. The hydration of catalytic bases involved in (partially rate-limiting) proton abstraction plays an important role not only in BLs, but also in other enzymes, such as ketosteroid isomerase and triosephosphate isomerase ^66, 67^, with reduced hydration of the base lowering the activation energy. Importantly, the barriers for deacylation from QM/MM reaction simulations obtained here agree very well with rates obtained experimentally, *if* OXA-163 and -405 do not adopt the reactive orientation of the DW (donating an H-bond to the carbapenem 6*α*-hydroxyethyl group). We show that adopting this arrangement in these variants, which lack carbapenemase activity, is indeed associated with an energy penalty, and how this is directly related to the changes in the *β*_5_-*β*_6_ loop. Specifically, changes to a water-mediated H-bond network between the carbapenem 6*α*-hydroxyethyl group and Thr213 in the *β*_5_-*β*_6_ loop, previously observed in crystal structures of OXA-163 and OXA-48 ^21, 24^, cause the different preference in DW orientation and H-bonding.

Whilst OXA-517 has been reported to have approximately the same turnover number for imipenem as OXA-48 (indicating efficient imipenem deacylation), its overall efficiency for imipenem breakdown is poor, due to a much increased *K*_M_ ^41^. Here, we show that this difference is probably related to a lower binding affinity for intact imipenem in OXA-517, in turn due to a subtle shift of the imipenem binding position, which impairs its interactions with the active site. No such differences in non-covalent binding were observed for OXA-163 and -405.

In summary, by using multiscale simulations, we conclude that the higher *k*_cat_ for imipenem breakdown by OXA-48 and -517, compared to OXA-163 and -405, can be attributed to the preference for different H-bonding interactions involving DW. Variations in the *β*_5_-*β*_6_ loop, mainly involving Thr213, result in distinct H-bond networks around the 6*α*-hydroxyethyl group of imipenem. This strong water-mediated H-bond network in OXA-48 leads to a preference for the DW H-bonding pattern favoured for reaction. In OXA-517, this H-bond network is weakened, but it can still stabilise the H-bonding pattern linked to high activity.

These insights from simulations complement and extend the information available through protein crystallography and kinetic studies. In particular, we provide a detailed explanation of how and why changes in the *β*_5_-*β*_6_ loop lead to a reduction in carbapenemase activity, something that was not yet understood. Our work thereby highlights how mutations can lead to very subtle effects, at the level of individual hydrogen bonds, which in turn lead to significant effects on enzyme activity. We thereby provide deeper understanding of how mutations around the active site can lead to modifications of the hydrolytic profile of *β*-lactamases, and thus also how such bacterial enzymes may adapt to confer resistance against *β*-lactam antibiotics. Such knowledge is important, because in general, clinical resistance mechanisms often arise from similarly small, structurally cryptic changes that collectively erode antibiotic efficacy. Furthermore, understanding exactly how loop modifications and (thereby) hydrogen bonding networks can influence *β*-lactam turnover may create opportunities for further rational design, e.g. new antibiotics that disrupt the key interactions required for efficient deacylation that we identify. Our results adds to similar insights obtained for other serine BL-substrate combinations, encompassing both BL-antibiotic ^22, 44, 68^ and BL-inhibitor systems ^69^, and thereby demonstrates how (multiscale) simulations can provide key insights related to understanding, and potentially combating, BL-conferred antibiotic resistance.

## Supporting information

Detailed description of applied restraints and the definition of the QM region used in the reaction simulations; multiple sequence alignment of OXA-48 and its variants; visualization of different imipenem 6*α*-hydroxyethyl orientations; full results of free energy surfaces for deacylation in different H-bond patterns and hydration states; cluster analysis of acylenzyme and Michaelis complex MM MD simulations; active site water and principal component analysis of acylenzyme MM MD simulations; salt bridge and Thr213 rotamer sampling of acylenzyme MM MD simulations; visualization of structures used in QM/MM potential energy calculations; energy decomposition of the difference in MM/GBSA binding energies (PDF)

## Data Availability

The input files, including the settings and initial structures for all simulations and QM/MM potential energy calculations, as well as the parameters for the carboxylated Lys73 and imipenem in both the free-ligand and Ser70-bound form, are made available at Zenodo: https://doi.org/10.5281/zenodo.18300786.

## Author contributions

DW and MvdK conceived the work. DW performed all simulations. DW and MvdK performed analysis and interpretation of results. MvdK supervised DW, with contributions from AJM and JS. DW and MvdK wrote the manuscript draft, with revisions and approval by all authors.

## Acknowledgments

DW and MWvdK thank the GW4 BIOMED2 DTP for DW’s studentship, grant MR/W006308/1 awarded to the Universities of Bath, Bristol, Cardiff and Exeter from the Medical Research Council (MRC)/UKRI. JS acknowledges funding from the U.K. Biotechnology and Biological Sciences Research Council (BBSRC, BB/W001187/1). AJM and JS acknowledge funding from the European Research Council under the European Horizon 2020 research and innovation program (PREDACTED Advanced Grant Agreement no. 101021207). All simulations and QM/MM potential energy calculations conducted using the facilities of the Advanced Computing Research Centre at the University of Bristol (http://www.bris.ac.uk/acrc/).

