## Supporting Information for "Hydrogen-bonding changes cause differences in imipenem breakdown activity in OXA-48 variants"

This Supporting Information contains:

- Detailed description of applied restraints and the definition of the QM region used in the reaction simulations (Note S1, Figure S3);
- Multiple sequence alignment of OXA-48 and its variants (Figure S1);
- Visualization of different imipenem 6$\alpha$-hydroxyethyl orientations (Figure S2);
- Experimental kinetic parameters for OXA-48 and its variants (Table S1);
- Full results of FESs in different H-bond patterns and hydration states and representative structures (Figures S4-S7; Table S2);
- Clustering, principal component, and active site water and analysis of acylenzyme MM MD simulations (Figure S8-S9; Table S3, Table S4);
- Salt bridge, flexibility changes (ΔRMSF) and Thr213 rotamer sampling of acylenzyme MM MD simulations (Figure S10-S13);
- Visualisation of X-Pro217 peptide bonds in X-ray crystal structures (Figure S14);
- Energetic quantification and visualization of structures from QM/MM potential energy calculations of hydrogen bonding networks (Table S5, Figure S15);
- Clustering of Michaelis complex MM MD simulations (Figure S16);
- Energy decomposition of the difference in MM/GBSA binding energies (Figure S17).

**Note S1**

To avoid changes in hydrogen bonding of the water surrounding the carboxylated Lys73 during the reaction simulations, one-sided harmonic distance and bond-angle restraints were applied between the KCX@OQ1 atom and the solvent water molecule donating a H-bond to it. Typically, KCX@OQ1 forms two H-bonds – one with the deacylating water and the other with this solvent water. The restraints came into force when the distance between the KCX@OQ1 atom and the solvent water hydrogen was above 2.2 Å, or when the bond angle of KCX@OQ1 (acceptor)-hydrogen-donor was below 150°. These restraints were introduced to prevent this solvent water from forming a hydrogen bond with KCX@OQ2 atom and thus altering its hydration state. In addition, a one-sided harmonic distance restraint was applied between the Ser70 hydroxyl oxygen and the electrophilic carbon of imipenem to maintain the covalently bound form of imipenem, permitting free sampling for distances below 1.6 Å. Depends on the DW H-bonding pattern, an additional one-sided harmonic distance restraint was also applied to maintain the corresponding DW H-bonding pattern. When the DW acts as H-bond donor to the imipenem hydroxyl group, the restraint is activated only if the distance between the hydroxyl hydrogen and the oxygen of a water molecule which also forms a H-bond with Try211 backbone oxygen exceeds 2.2 Å. While when the DW acts as H-bond acceptor to the imipenem hydroxy group, the restraint is applied between the imipenem hydroxyl group hydrogen and the oxygen of the DW, allowing free sampling when the distance is below 1.9 Å. Force constants for all restraints and reaction coordinates were 100 kcal $mol^{-1}$ $Å^{-2}$ (distance type restraints) or 100 kcal $mol^{-1}$ $rad^{-2}$ (bond angle type restraints).

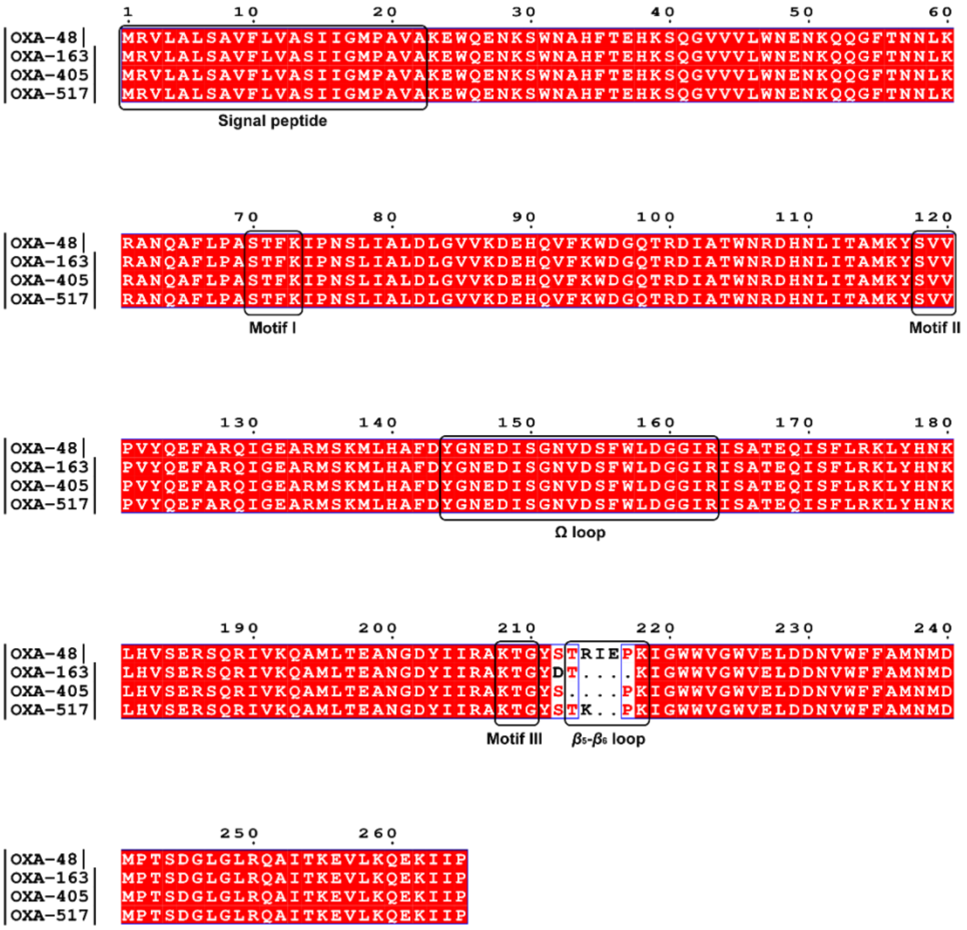

**Figure S1.** Multiple sequence alignment of OXA-48-like proteins performed with NCBI BLAST [1] and mapped using the ESPrint webserver [2]. The signal peptide, Ω loop, $\beta_5$-$\beta_6$ loop and three conserved motifs are boxed. GeneBank IDs for enzymes are: LN864820.1 for OXA-48; HQ700343.1 for OXA-163; KM589641.1 for OXA-405 and KU878974.1 for OXA-517.

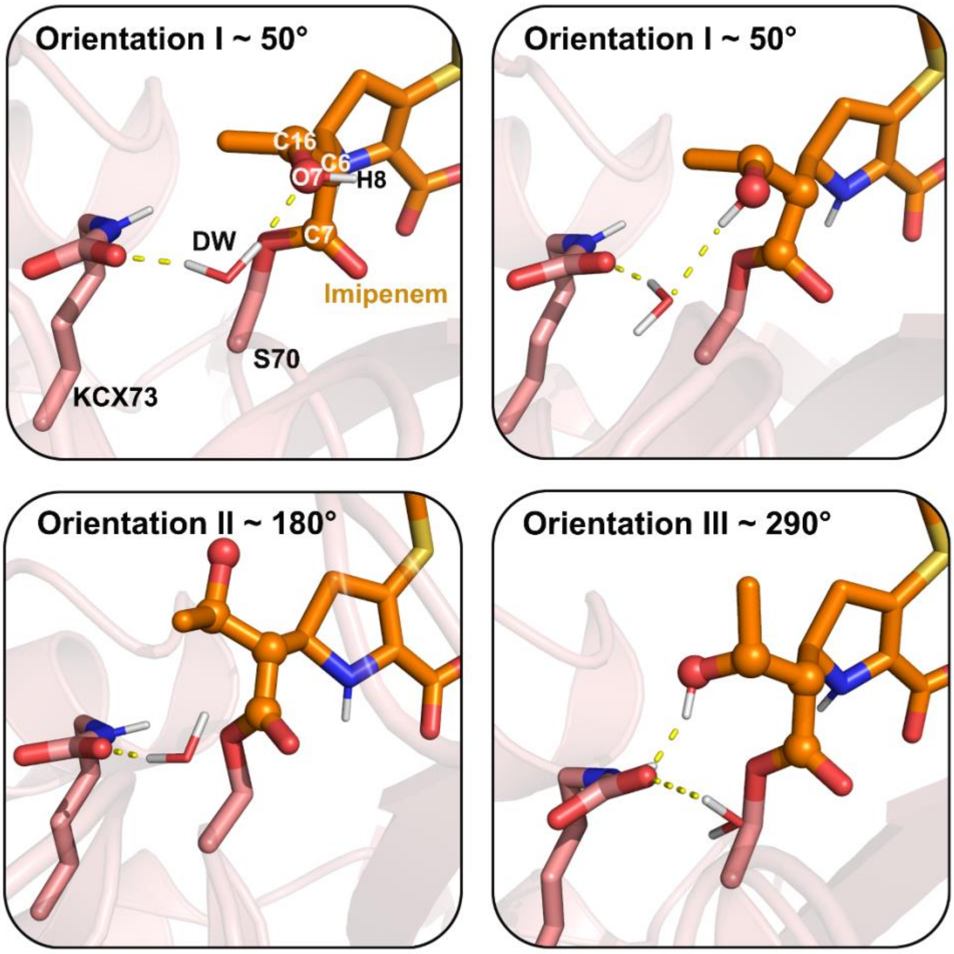


**Figure S2.** The imipenem 6α-hydroxyethyl can adopt three orientations, dihedral angle around 50° (orientation I), 180° (orientation II), and 290° (orientation III). The atoms used to determine the dihedral angle are shown as spheres. In orientation I, the deacylating water forms two different H-bonding patterns with the 6α-hydroxyethyl hydroxyl group. Only polar hydrogens are shown. The sidechains of carboxylated Lys73 (pink) and imipenem (orange) are shown as sticks. H-bonds are shown as yellow dash lines.

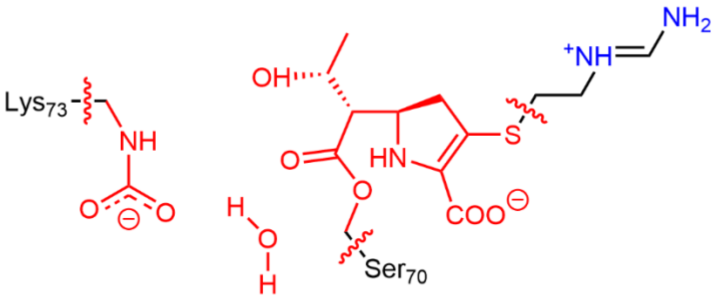


**Figure S3.** QM region in QM/MM MD reaction simulations. The atoms of QM region are coloured in red, and the breaks between QM and MM region are shown with wavy lines.

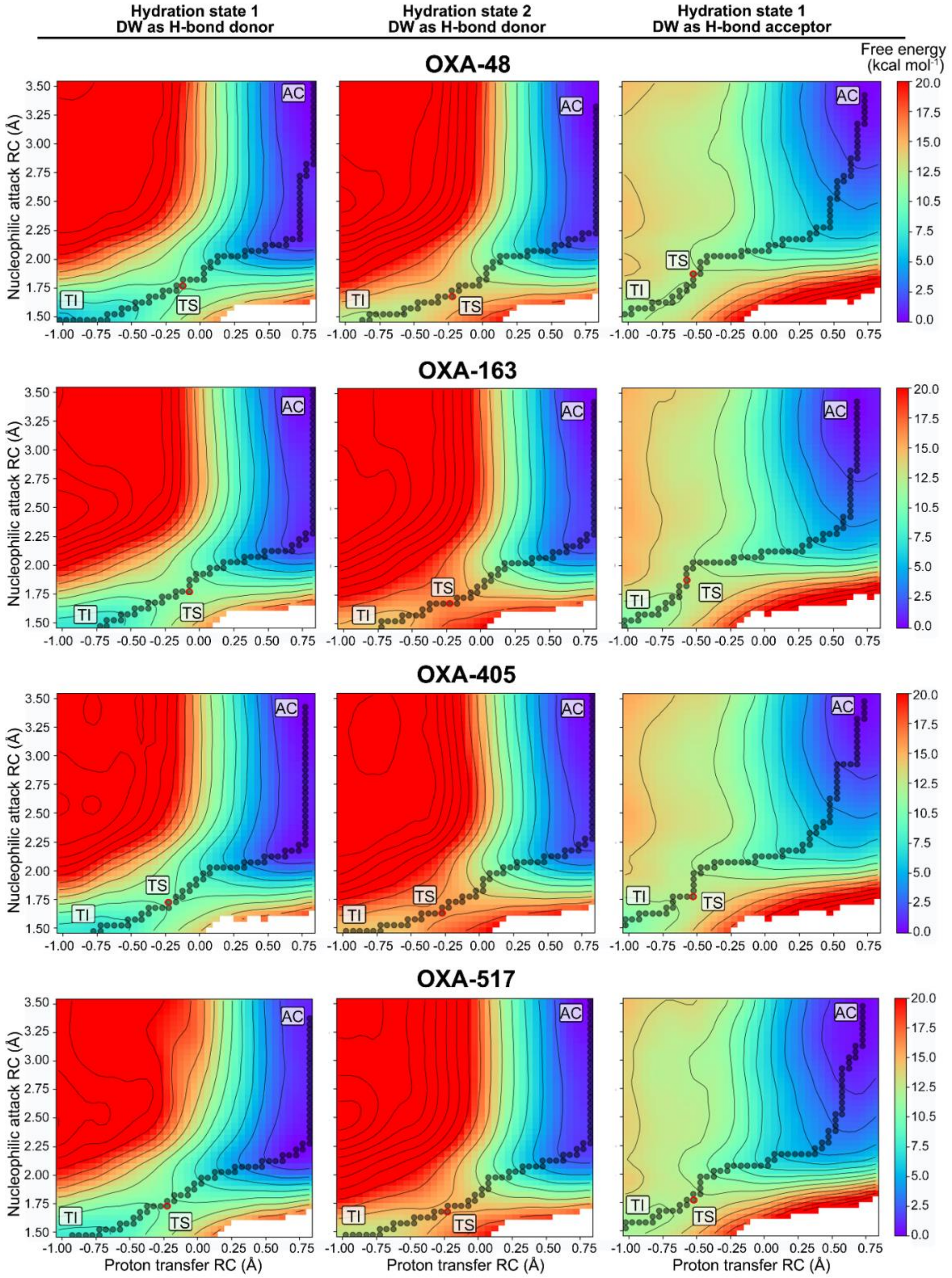


**Figure S4.** Free energy surfaces from DFTB2/ff14SB simulations of imipenem deacylation by OXA-48-like proteins, with minimum free energy paths (MFEPs) indicated by circles. Surfaces correspond to the lowest calculated barriers among three independent QM/MM umbrella sampling runs for each of the three different hydration states and H-bonding patterns indicated. AC = acylenzyme, TS = transition state, TI = tetrahedral intermediate.

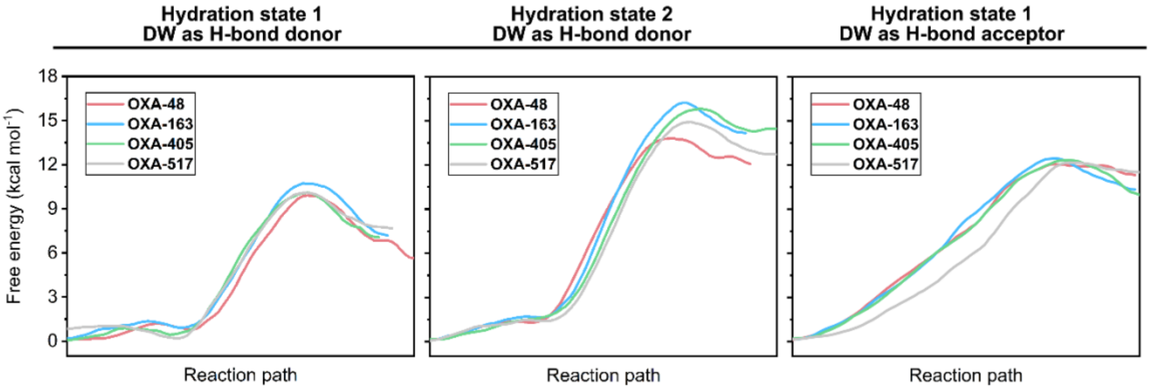


**Figure S5.** Free energy profiles from DFTB2/ff14SB simulations of imipenem deacylation by OXA-48-like proteins. Profiles correspond to the minimum free energy paths of the surfaces shown in Figure S3.

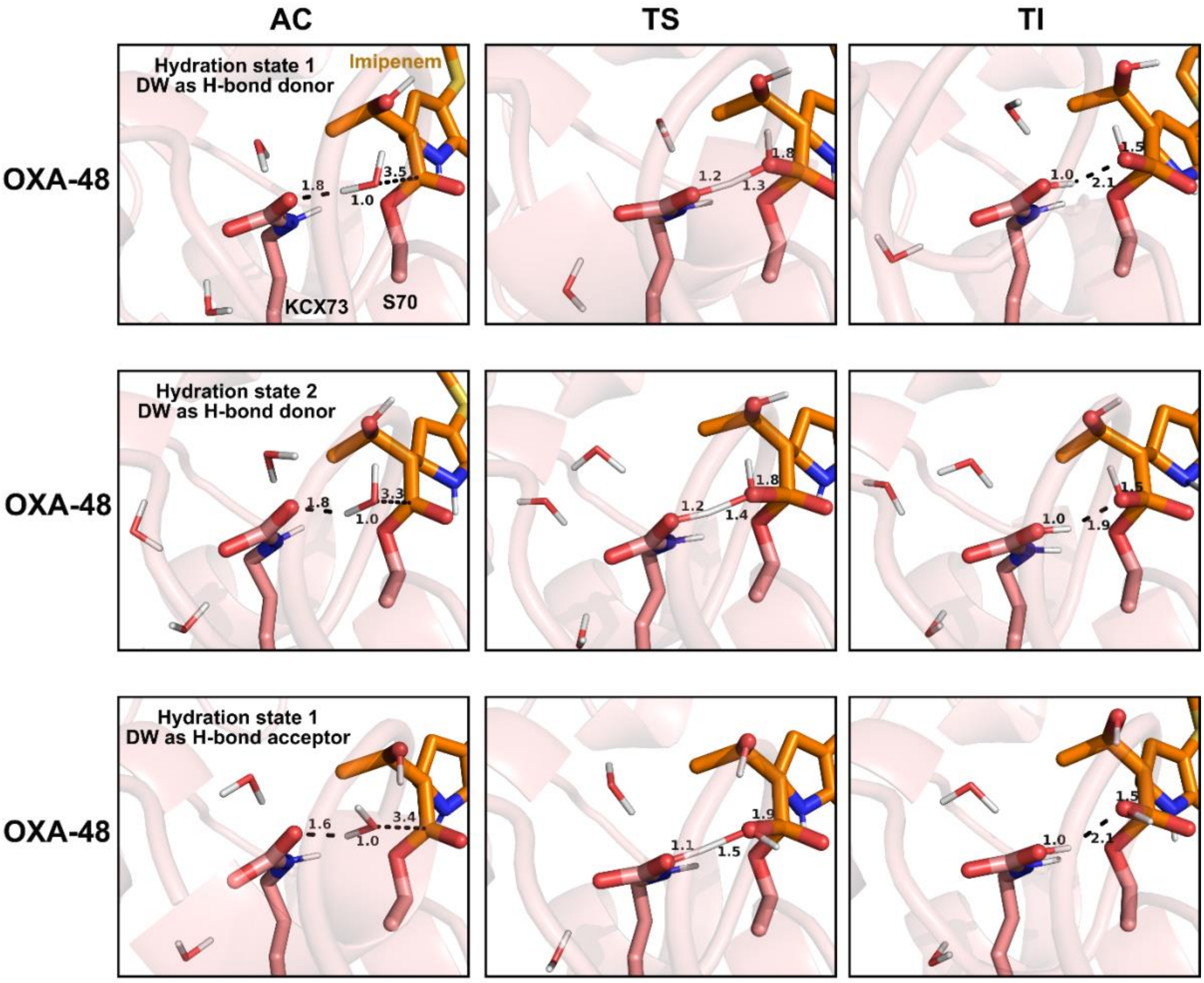


**Figure S6.** Representative acylenzyme, transition state and tetrahedral intermediate structures from DFTB2/ff14SB simulations of imipenem deacylation by OXA-48. AC = acylenzyme, TS = transition state, TI = tetrahedral intermediate. Structures were obtained from the corresponding free energy surfaces shown in Figure S3. Only polar hydrogens are shown. Key residues and imipenem (orange) are shown as sticks, and the reaction coordinates are labelled in Å.

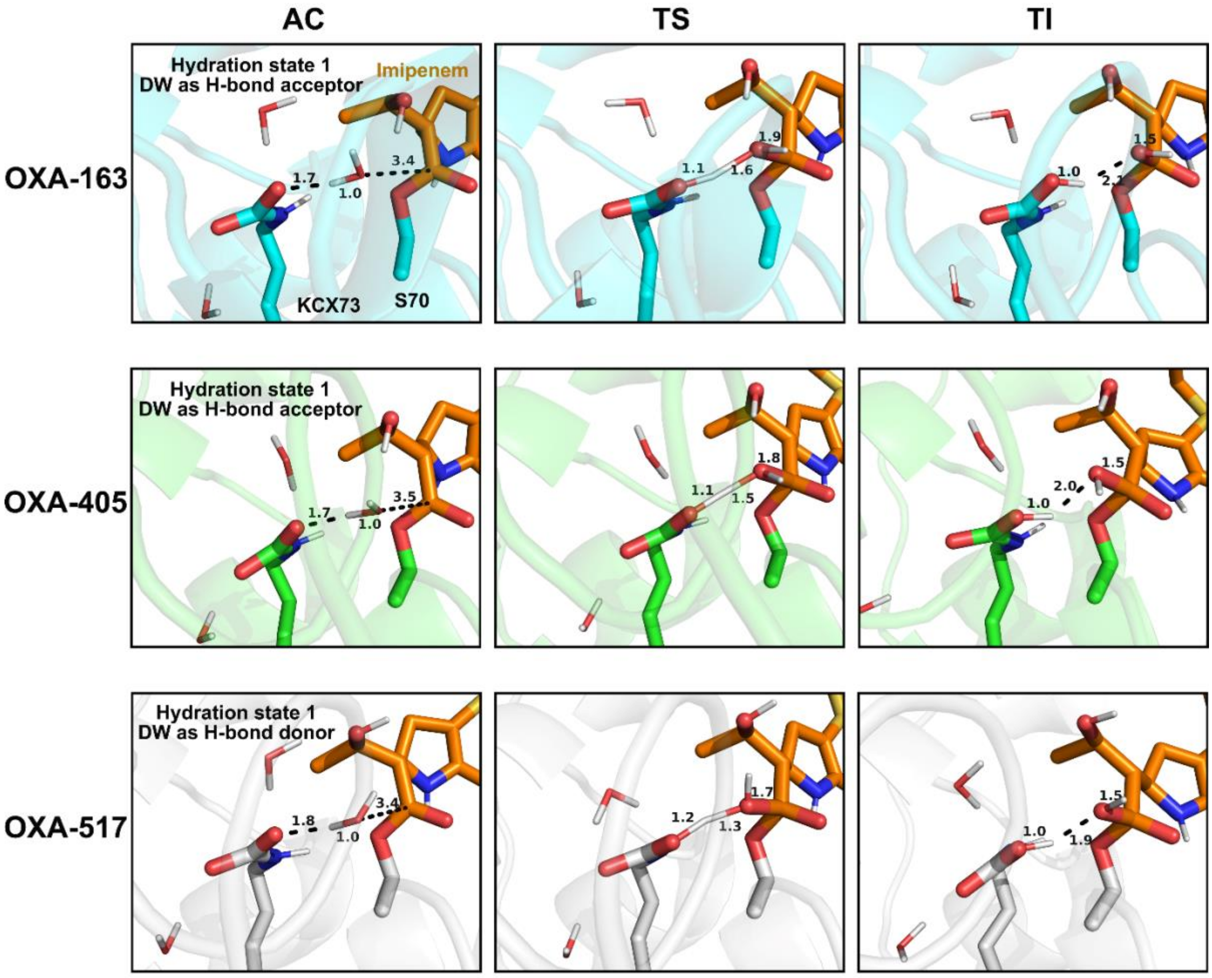


**Figure S7.** Representative acylenzyme, transition state and tetrahedral intermediate structures from DFTB2/ff14SB simulations of imipenem deacylation by OXA-163, -405 and -517. AC = acylenzyme, TS = transition state, TI = tetrahedral intermediate. Only the hydration state and H-bonding patterns that used for the fitted line in Figure 3B are considered. Structures were obtained from the corresponding free energy surfaces shown in Figure S3. Only polar hydrogens are shown. Key residues and imipenem (orange) are shown as sticks, and the reaction coordinates are labelled in Å.

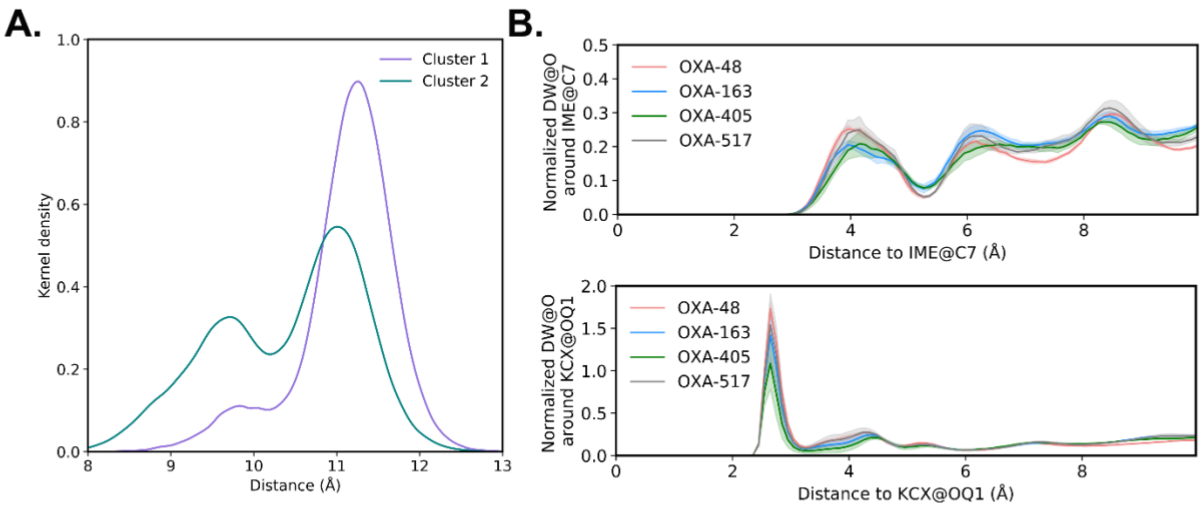


**Figure S8.** (A) Distribution (kernel-density estimate) of the distances between C$\alpha$ atoms of Val120 and Leu158 across the two clusters identified by clustering on C$\alpha$ atom RMSD of all acylenzyme MM MD simulations of the variants combined. (B) Radial distribution function of water molecules around selected atoms in the acylenzyme MM MD simulations. The 95% confidence interval is shown as a shaded region, and was calculated as $\bar{x} \pm 1.96\sigma/\sqrt{n}$, where $\bar{x}$ is the sample mean, $\sigma$ is the standard deviation, and $n$ is the number of independent MD simulations per system ($n$ = 5 for each system).

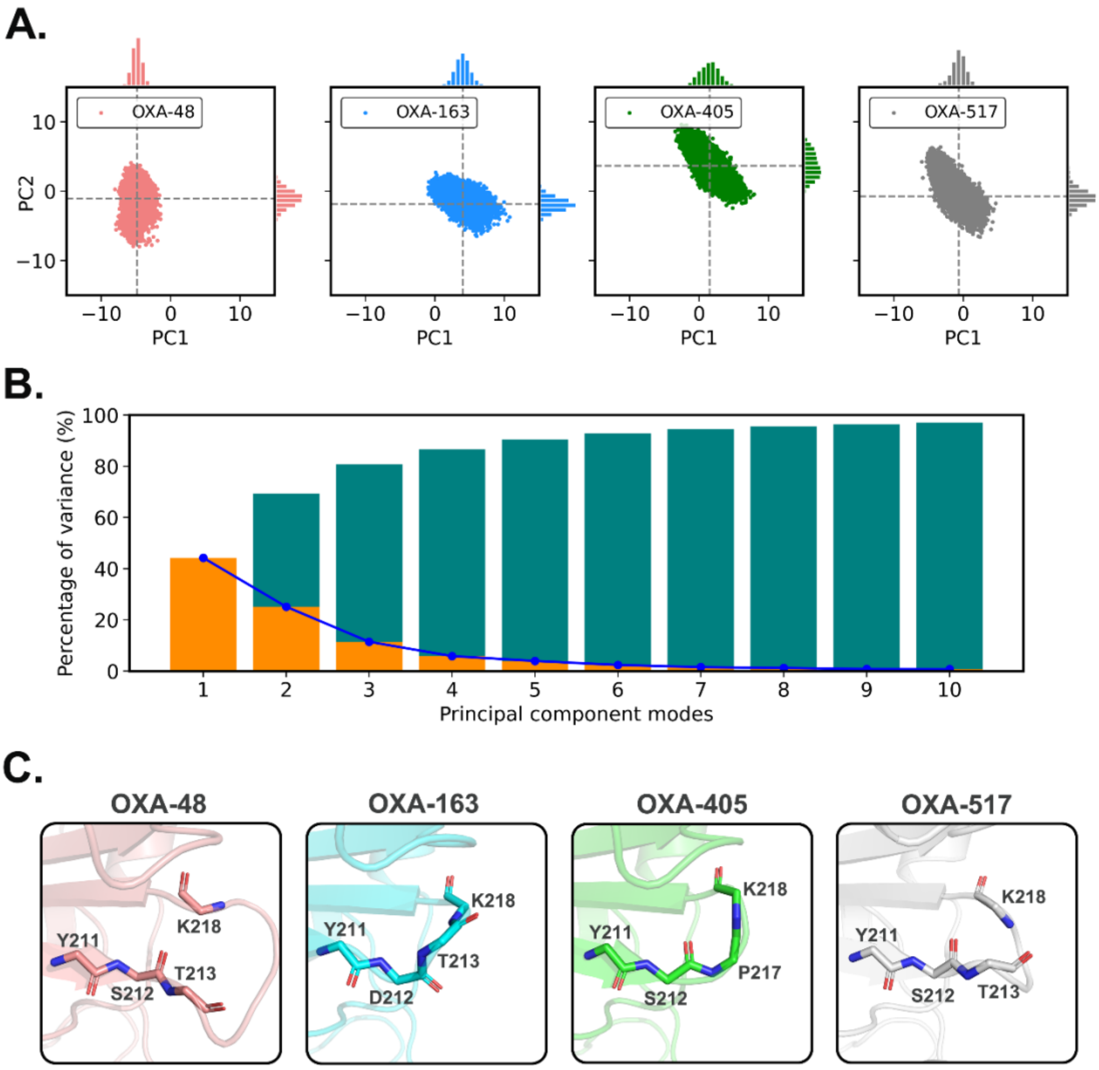


**Figure S9.** Principal component analysis of the $\beta_5$-$\beta_6$ loop backbone atoms (N, CA, C and O) based on all acylenzyme MM MD simulations combined. (A) Principal components (PC1 and PC2) values of backbone atoms of common $\beta_5$-$\beta_6$ loop residues, Tyr211 and Ser/Asp212. The mean number of each principal component is shown as black dash line. (B) The variance of first 10 principal components. The coloured bars show the individual (orange) and cumulative (green) variances. (C) Representative $\beta_5$-$\beta_6$ loop structures based on the average values of principal components.

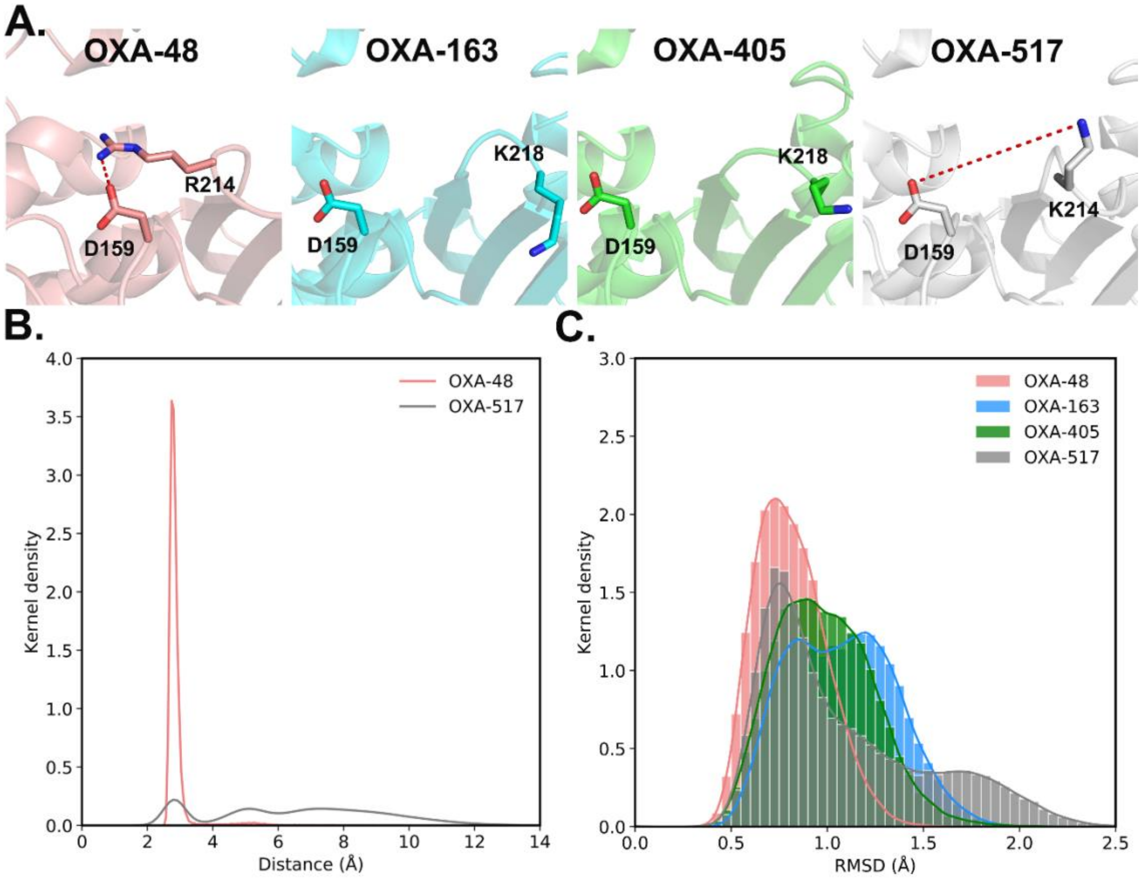


**Figure S10.** Salt bridge interaction between the Ω loop and the $\beta_5$-$\beta_6$ loop as observed in the combined acylenzyme MM MD simulations. (A) Structures of the salt bridge between Arg/Lys214 and Asp159 in different OXA-48-like proteins. (B) Distribution (kernel density estimate) of the closest distance sampled between the sidechain nitrogens of Arg/Lys214 and carboxylate oxygens of Asp159. (C) Histogram of Ω loop heavy atom RMSD values in different acylenzyme complexes. Trajectories were first aligned based on all C$\alpha$ atoms apart from the Ω loop. Bin width is 0.05 Å.

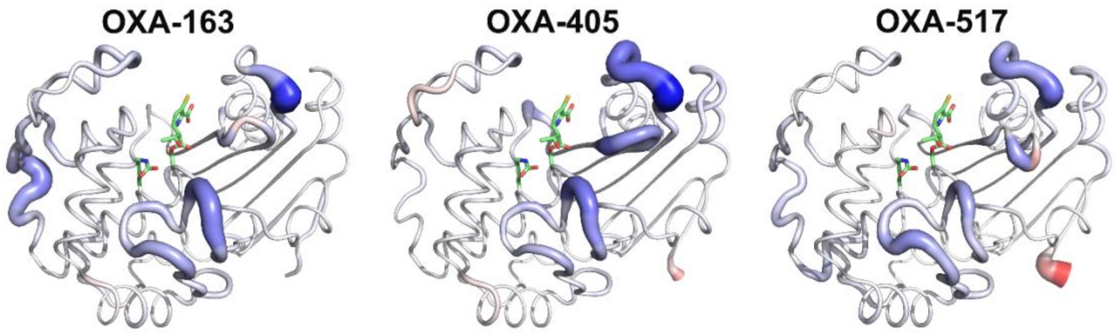


**Figure S11.** $\Delta RMSF_{C\alpha}$ (compared to OXA-48) calculated from the combined acylenzyme MM MD simulations, displayed on structures. The thickness and colour of the tube are determined by $\Delta RMSF_{C\alpha}$ values obtained from Figure 3C (blue indicating higher flexibility than OXA-48, red lower). The sidechain of the carboxylated Lys73 and the QM part of imipenem (without the C2 tail) are shown in sticks, for reference.

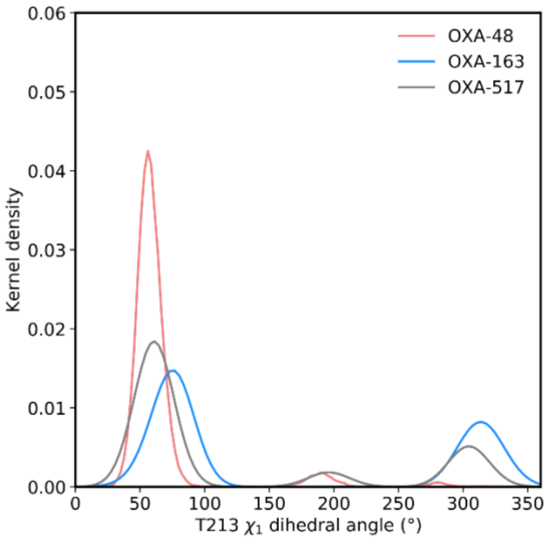


**Figure S12.** Distribution (kernel-density estimate) of the Thr213 $\chi1$ dihedral angle sampled in the combined acylenzyme MM MD simulations.

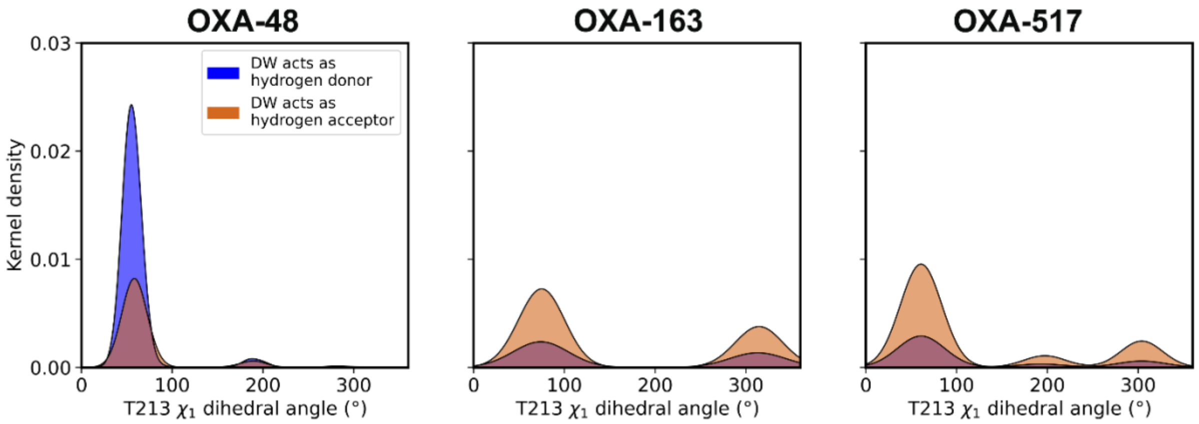


**Figure S13.** The Thr213 sidechain dihedral angle sampling (kernel-density estimates) in those frames of the acylenzyme MM MD simulations with a reactive DW position (≤ 3.0 Å between DW@O and KCX@OQ1 *and* ≤ 3.5 Å between DW@O and the electrophilic carbon of imipenem), separated by DW H-bonding patterns. (The area under the curve of each plot is 1, even though the systems have a different number of frames with the reactive DW position).

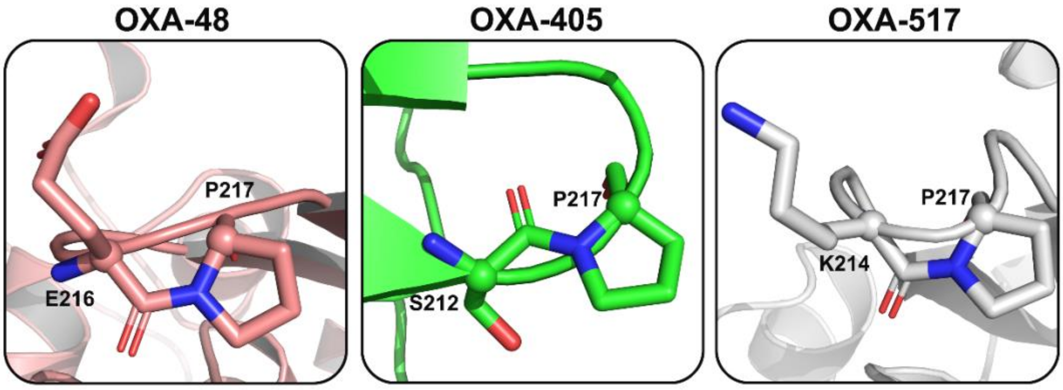


**Figure S14.** The X-Pro peptide bond in the $\beta_5$-$\beta_6$ loop as observed in X-ray crystal structures of OXA-48 (PDB ID: 6P97 [3], chain A), OXA-405 (PDB ID: 5FDH [4], chain A) and OXA-517 (PDB ID: 6HB8 [5], chain A). Residue X and Pro are shown as sticks, and the C$\alpha$ atoms are shown as spheres. In OXA-48 and OXA-517, the X-Pro peptide bond is in the cis conformation, while it is in the trans conformation in OXA-405. The X-Pro peptide bond remains in these conformations during all simulations. This proline is deleted in OXA-163, hence OXA-163 is not shown here.

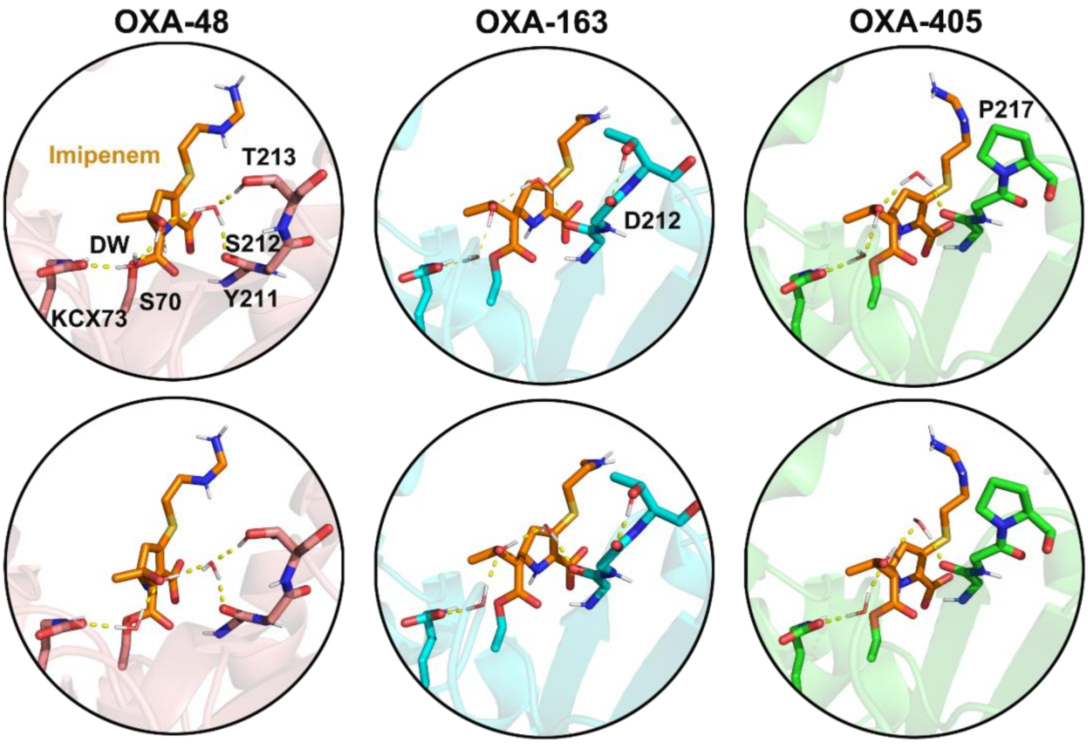


**Figure S15.** Representative structures at the local QM/MM potential energy minima obtained from scanning along the of dihedral angle of the imipenem 6α-hydroxyethyl hydroxyl group. Minima correspond to the two different deacylating water H-bonding patterns (top and bottom) in OXA-48 (left), OXA-163 (middle) and OXA-405 (right) acylenzyme structures. Only polar hydrogens are shown, for clarity. Key residues (grey) and imipenem (orange) are shown as sticks and H-bonds are shown as yellow dash lines.

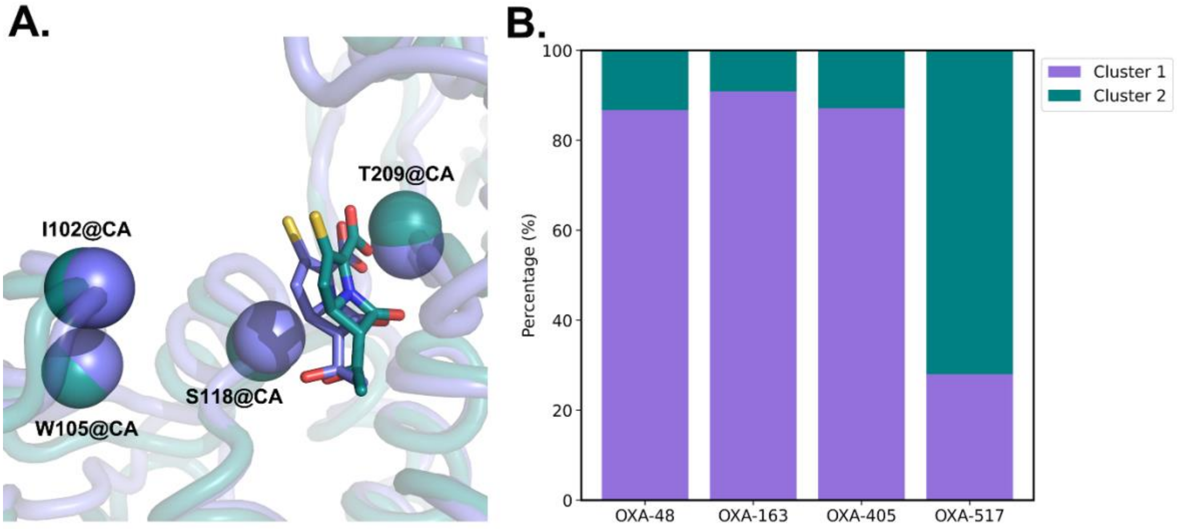


**Figure S16.** Results from clustering of the combined Michaelis complex MM MD simulations based on the RMSD of the imipenem core structure (after alignment on the active site). (A) Representative structures of the two identified clusters. Residues that show different positions between the clusters are shown as spheres. Only the core structure of imipenem (as used in clustering) is shown as sticks. (B) The percentage of frames belonging to each cluster per enzyme variant.

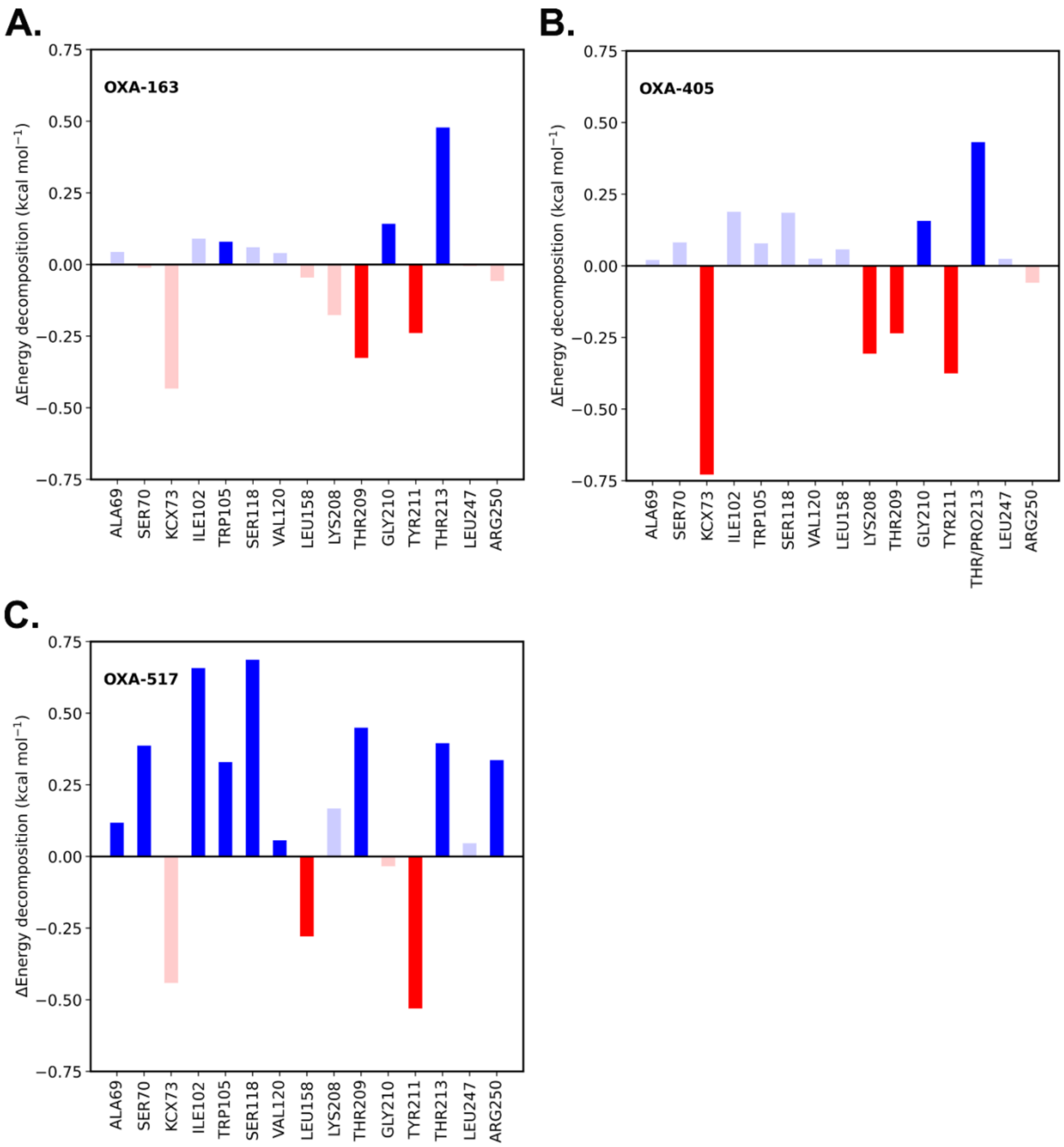


**Figure S17.** Per-residue decomposition of the difference in MM/GBSA binding energies (ΔEnergy) for OXA-163 (A), OXA-405 (B) and OXA-517 (C), compared to OXA-48. Residues numbers of OXA-48 are used here. Only residues which contribute to binding energy significantly are shown (absolute value of energy contribution >0.5 kcal mol$^{-1}$). Contributions increasing binding energy (decrease of affinity) w.r.t OXA-48 are positive. Differences in contributions shown in transparent bars are not statistically significant ($p > 0.05$).

**Table S1. Kinetic parameters of OXA-48 and its variants with imipenem**

| Enzyme | $k_{cat}$ ($s^{-1}$) | $K_M$ (μM) |
|---|---|---|
| OXA-48[a] | 5 | 13 / 3[c] |
| OXA-163[b] | 0.03 | 520 / 3.8[c] |
| OXA-405[d] | 0.1 | 532 |
| OXA-517[e] | 6 | 411 |

[a] Kinetic parameters for OXA-48 are from Docquier *et al* [6].

[b] Kinetic parameters for OXA-163 are from Oueslati *et al* [7].

[c] Different experimental $K_M$ values of OXA-48 and -163 are reported by Stojanoski *et al* [8].

[d] Kinetic parameters for OXA-405 are from Oueslati *et al* [4].

[e] Kinetic parameters for OXA-517 are from Dabos *et al* [5].

**Table S2. Calculated free energy barriers for imipenem deacylation**

| Enzyme | DW acts as hydrogen donor | | DW acts as hydrogen acceptor |
|---|---|---|---|
| | Hydration state 1 (kcal $mol^{-1}$)[a] | Hydration state 2 (kcal $mol^{-1}$)[a] | Hydration state 1 (kcal $mol^{-1}$)[a] |
| OXA-48 | 10.8 ± 0.8 | 14.0 ± 0.1 | 12.8 ± 0.7 |
| OXA-163 | 11.5 ± 0.7 | 16.5 ± 0.2 | 13.0 ± 1.0 |
| OXA-405 | 11.1 ± 0.9 | 15.9 ± 0.1 | 12.5 ± 0.1 |
| OXA-517 | 10.9 ± 0.7 | 15.4 ± 0.5 | 13.1 ± 0.8 |

[a] Standard deviations obtained from three independent US calculations.

**Table S3. Clustering of OXA-48-like proteins in acylenzyme complexes based on C*α* atoms**

| Enzyme | Cluster 1 (fraction) | Cluster 2 (fraction) |
|---|---|---|
| OXA-48 | 77.3% | 22.7% |
| OXA-163 | 30.1% | 69.9% |
| OXA-405 | 35.4% | 64.6% |
| OXA-517 | 66.2% | 33.8% |

**Table S4. Frequency of deacylating water in reactive position[a] in acylenzyme complexes**

| Enzyme | Fraction |
|---|---|
| OXA-48 | 9.2% |
| OXA-163 | 7.9% |
| OXA-405 | 5.7% |
| OXA-517 | 6.5% |

[a] Based on an 'ideal' DW position for deacylation of ≤ 3.0 Å between DW@O and KCX@OQ1 *and* ≤ 3.5 Å between DW@O and the electrophilic carbon of imipenem.

**Table S5. Potential energy differences for different H-bonding patterns of the deacylating water**

| Enzyme | DFTB2/ff14SB energy (kcal $mol^{-1}$) [a] | M06-2X/def2-TZVP//ff14SB energy (kcal $mol^{-1}$) [a, b] |
|---|---|---|
| OXA-48 | 2.9 ± 1.0 | 2.6 ± 0.6 |
| OXA-163 | 5.2 ± 0.8 | 5.3 ± 0.8 |
| OXA-405 | 8.6 ± 1.5 | 7.5 ± 1.0 |
| OXA-517 | 1.1 ± 1.4 | 2.2 ± 0.4 |

[a] Standard deviations obtained from three independent calculations.

[b] Calculated by replacing only the QM region DFTB2 energy with the M06-2X/def2-TZVP energy.
